# A regression based approach to phylogenetic reconstruction from multi-sample bulk DNA sequencing of tumors

**DOI:** 10.1101/2024.04.23.590844

**Authors:** Henri Schmidt, Benjamin J. Raphael

## Abstract

**Motivation:** DNA sequencing of multiple bulk samples from a tumor provides the opportunity to investigate tumor heterogeneity and reconstruct a phylogeny of a patient’s cancer. However, since bulk DNA sequencing of tumor tissue measures thousands of cells from a heterogeneous mixture of distinct sub-populations, accurate reconstruction of the tumor phylogeny requires simultaneous deconvolution of cancer clones and inference of ancestral relationships, leading to a challenging computational problem. Many existing methods for phylogenetic reconstruction from bulk sequencing data do not scale to large datasets, such as recent datasets containing upwards of ninety samples with dozens of distinct sub-populations.

**Results:** We develop an approach to reconstruct phylogenetic trees from multi-sample bulk DNA sequencing data by separating the reconstruction problem into two parts: a structured regression problem for a fixed tree 𝒯, and an optimization over tree space. We derive an algorithm for the regression sub-problem by exploiting the unique, combinatorial structure of the matrices appearing within the problem. This algorithm has both asymptotic and empirical improvements over linear programming (LP) approaches to the problem. Using our algorithm for this regression sub-problem, we develop *fastBE*, a simple method for phylogenetic inference from multi-sample bulk DNA sequencing data. We demonstrate on simulated data with hundreds of samples and upwards of a thousand distinct sub-populations that *fastBE* outperforms existing approaches in terms of reconstruction accuracy, sample efficiency, and runtime. Owing to its scalability, *fastBE* also enables phylogenetic reconstruction directly from indvidual mutations without requiring the clustering of mutations into clones. On real data from fourteen B-progenitor acute lymphoblastic leukemia patients, *fastBE* infers similar phylogenies to the existing, state-of-the-art method, but with fewer violations of a widely used evolutionary constraint and better agreement to the observed mutational frequencies. Finally, we show that on two patient-derived colorectal cancer models, *fastBE* also infers phylogenies with less violation of a widely used evolutionary constraint compared to existing methods, and leads to distinct interpretations of the intra-tumor heterogeneity.

**Availability:** *fastBE* is implemented in C^**++**^ and is available at: github.com/raphael-group/fastBE.

## 1 Introduction

Tumor evolution is characterized by the accumulation of somatic genomic alterations that alter the fitness of sub-populations of cells, leading to unregulated growth. Over the past ten years, high-coverage DNA sequencing of bulk tumor samples has proven tremendously successful in deciphering this complex evolutionary process [1–3]. There are now dozens of computational techniques [4–12] to accurately and efficiently identify distinct sub-populations of cells in a tumor sample and reconstruct the evolutionary history, or a phylogeny, of these populations. Application of these techniques can help identify the genomic alterations that drive tumor growth [13, 14]. Recent studies have demonstrated that intra-tumor heterogeneity is more prevalent than previously reported (reviewed in [15]). For example, the TracerX Consortium [2] found up to fifteen distinct sub-populations of cells, or *subclones*, from multi-region sequencing of a patient biopsy of non-small-cell lung cancer. Further, they noticed that without this multi-region sequencing, up to 65% of subclones and 76% of subclonal mutations would have been missed, suggesting that perhaps further heterogeneity could be uncovered by increasing the number of sequenced samples. In tandem with decreasing sequencing costs, high levels of intra-tumor heterogeneity have led to bulk-DNA sequencing datasets with an increasingly large number of samples and subclones – containing up to 90 samples and 26 subclones for a single cancer patient [16].

While numerous methods have been developed to build phylogenies from bulk DNA sequencing of tumors, few of these methods scale to datasets with dozens of samples and subclones from the same patient. Recently [11] demonstrated that existing methods fail to scale beyond even ten subclones, making their application to datasets with high amounts of intra-tumor heterogeneity challenging. As another consequence of poor scalability, all existing methods – except the newly introduced method Orchard [12] – infer phylogenies using summary statistics computed from *clusters of mutations* rather than the individual mutation read counts, potentially missing valuable phylogenetic signal.

### Contribution

In this work, we describe a *structured regression* formulation of the successful matrix factorization model [5–7, 9, 10] for phylogenetic inference from somatic single nucleotide variants (SNVs) measured via DNA sequencing of one or more bulk tumor samples from the same patient. In particular, we identify a tractable, ℓ_1_-regression problem hidden within the NP-complete, variant allele frequency (VAF) factorization problem [5, 17] – analogous to the *method of minimum evolution* in species phylogenetics [18, 19] – where a tractable regression sub-problem [20] is solved within an NP-complete [21, 22] optimization problem. By studying the unique, combinatorial *structure* of the clonal matrices [5] appearing within this ℓ_1_-regression sub-problem, we derive an algorithm which obtains both asymptotic and empirical improvements over a naїve, linear programming based approach. Further, our regression algorithm efficiently recomputes the solution to the ℓ_1_-regression sub-problem upon slight modifications to the tree topology, such as the addition of vertices and subtree prune-and-regraft (SPR) operations [23].

Utilizing our fast ℓ_1_-regression algorithm and incorporating recently introduced combinatorial search techniques [12], we develop a simple method, *fastBE* (*fast B*ulk *E*volution), for phylogenetic inference from multi-sample bulk DNA sequencing data. We show that on simulated data, *fastBE* outperforms existing methods for inferring the ground truth phylogeny across a variety of metrics – including sample efficiency – while running orders of magnitude faster. For example, *fastBE* solves simulated instances with up to 1000 clones and 100 samples in under half an hour. Applying *fastBE* to a multi-sample dataset from fourteen patients with acute lymphoblastic leukemias [16], we show that our inferred phylogenies are similar to those inferred by Pairtree [11], but better recapitulate observed mutational frequencies and possess fewer violations of the *sum condition* [5–7, 24]. Finally, we demonstrate that on two patient-derived colorectal cancer models *fastBE* finds phylogenies with fewer violations of the sum condition compared to existing methods.

## 2 Background and Related Work

Following previous work, we restrict attention to somatic, single-nucleotide mutations in copy neutral regions as phylogenetic characters of cancer evolution. We assume that each *genomic locus* is mutated exactly once during this evolutionary process, known as the *infinite sites assumption* [5–11]. The problem of inferring a phylogeny from DNA sequencing data from multiple bulk samples then corresponds to a matrix factorization problem – called the variant allele frequency factorization problem (VAFFP) [5] - which we describe below.

Under these assumptions, the evolutionary history of a tumor is described as a rooted, phylogenetic tree 𝒯= (*V*(𝒯), *E*(𝒯) where the vertices *V* correspond to sub-populations of cells containing identical sets of mutations, or *clones*, and the edges *E* define ancestral relationships between the clones. Mathematically, a mutation is represented by its position *j* in the genome and a clone *b*_*i*_ is a length *n* binary vector where *b*_*i,j*_ = 1 (resp. *b*_*i,j*_ = 0) denotes the presence (resp. absence) of mutation *j* in the *i*^th^ clone^1^. As each mutation occurs exactly once during the evolutionary process, there are *n* distinct clones *b*_*i*_, and they form an important subclass of *perfect phylogenies* [28, 29] where the internal vertices, in addition to the leaves, are labeled [5, 30]. Then, the evolutionary history of a tumor is given by a vertex labeled, perfect phylogeny with *n* vertices, which we call an *n-clonal tree*^2^ to emphasize that each of the *n* vertices correspond to a distinct tumor clone.

### Definition 1.

*A rooted tree 𝒯* =( *V, E*) *with vertices V* = [*n*] *is an n-clonal tree if each edge* (*i, j*) *is labeled by mutation j*. *We simply write clonal tree when n is clear by context*.

The root *r*(𝒯) of an *n*-clonal tree 𝒯 is assigned the unique mutation that does not appear as a label on any edge and is denoted *r* when the tree is clear by context. The vertices and edges of 𝒯 are denoted as *V*(𝒯) and *E*(𝒯). The parent of a non-root, vertex *i* in *V*(𝒯) is written as *δ* (*i*) and the set of children of a vertex *i* is written as *C* (*i*). The depth *d*(*i*) of a vertex *i* in *V*(𝒯) is the number of edges on the path from *r*(𝒯) to *i*. The depth of the tree 𝒯 is the maximum depth of any vertex *i* in *V*(𝒯). The clone *b*_*i*_ corresponds to vertex *i* of the *n*-clonal tree and contains all mutations occurring on the unique path from *r* to *i* in 𝒯. We summarize all clones in 𝒯 with an *n-clonal matrix* where each clone is a row of the matrix.

### Definition 2.

*The n-clonal matrix B*_𝒯_ = [*b*_*i,j*_] *is the n-by-n binary matrix such that b*_*i,j*_ = 1 *if and only if either i is a descendant of j in* 𝒯 *or i* = *j*. *We drop the subscript* 𝒯 *when it is clear by context*.

The model of bulk DNA sequencing is then as follows: each of *m* samples consists of a mixture of distinct clones and the sequencing experiment measures the frequency of all mutations in this mixture. More formally, it is assumed that each of the *m* measurements *f*_*i*_ ∈ ℝ^*n*^ are *convex combinations* [31] of the *n* clones in 𝒯. That is,

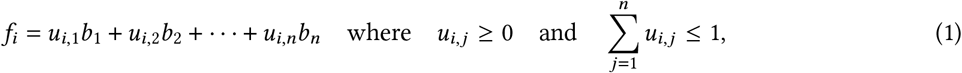

where clone *b*_*i*_ is the *i*^th^ row of *B*. This model is often summarized compactly in matrix notation as *F* = *UB*, where *F* = [*f*_*i*_]and *U* = [*u*_*i*_] are *m*-by-*n* matrices, *f*_*i,j*_ is the frequency of mutation *j* in sample *i*, and *u*_*i,j*_ is the fraction of clone *j* in sample *i*. As such, we call *F* a *frequency matrix* and any right stochastic matrix *U* a *usage matrix*.

Under this model, the problem of reconstructing the evolutionary history of the tumor becomes equivalent to factoring the observed frequency matrix *F* into its constituents *U* and *B*. While it would be desirable to solve this factorization problem exactly, imperfect measurement makes this challenging in practice and instead, most methods attempt to infer *U* and *B* such that *F* ≈ *U B*. For example, CITUP [7] attempts to find *U* and *B* such that 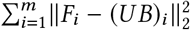 is minimized. In the general setting, we have a loss function *L* (*F, U, B*) that provides a measure of error between the observed frequency matrix *F* and the inferred matrices *U* and *B*.

### Problem 1

(The Variant Allele Frequency *L*-Factorization Problem (*L*-VAFFP)). *Given a frequency matrix F, find a clonal matrix B and a usage matrix U such that the loss L*(*F, U, B*) *is minimized*.

Multiple variations of the *L*-VAFFP have been studied in the literature, with different choices of loss function *L*(*F, U, B*) and additional constraints, as summarized in Table 1. In particular, CITUP [7] jointly clusters mutations and infers a phylogeny, using a regularized *L*_2_ loss to avoid overfitting on the number of mutation clusters. CITUP is an exact algorithm to solve this problem based off exhaustive enumeration of clonal trees and quadratic integer programming. LICHeE [6] also clusters mutations, and minimizes the total squared violation of the *sum condition* [5–7, 24] in their inferred tree 𝒯 using quadratic programming. AncesTree [5] on the other hand, does not cluster mutations, and studies two different loss functions. The first loss they study is the 0-1 loss *L*(*F, U, B*) = 𝟙 (*F* ≠ *U B*), which is zero if and only if *F* = *U B*. Interestingly, they show that under this loss function, the variant allele frequency factorization problem is NP-complete, which implies that it is also NP-complete under any *L*_*p*_ loss. As real data is quite noisy, they also study a variant of the *L*_1_ loss obtained from adding the hard constraint that |*F*_*i j*_ *UB*_*ij*_ |≤ *ϵ*_*i,j*_ for some *ϵ >* 0. For both loss functions, they use an integer linear programming formulation to solve the problem exactly. CALDER [10] builds off of the approach of AncesTree, but adds an additional hard constraint that the inferred matrices *U* and *B* are *longitudinally consistent*, leveraging the temporal information present in certain experimental settings. Further, rather than minimizing the *L*_1_ loss, they minimize the *L*_0_ loss on the usage matrix *U*. Again, they utilize integer linear programming to solve this optimization problem exactly. A complete description of all methods is summarized compactly in Table 1.

**Table 1:**
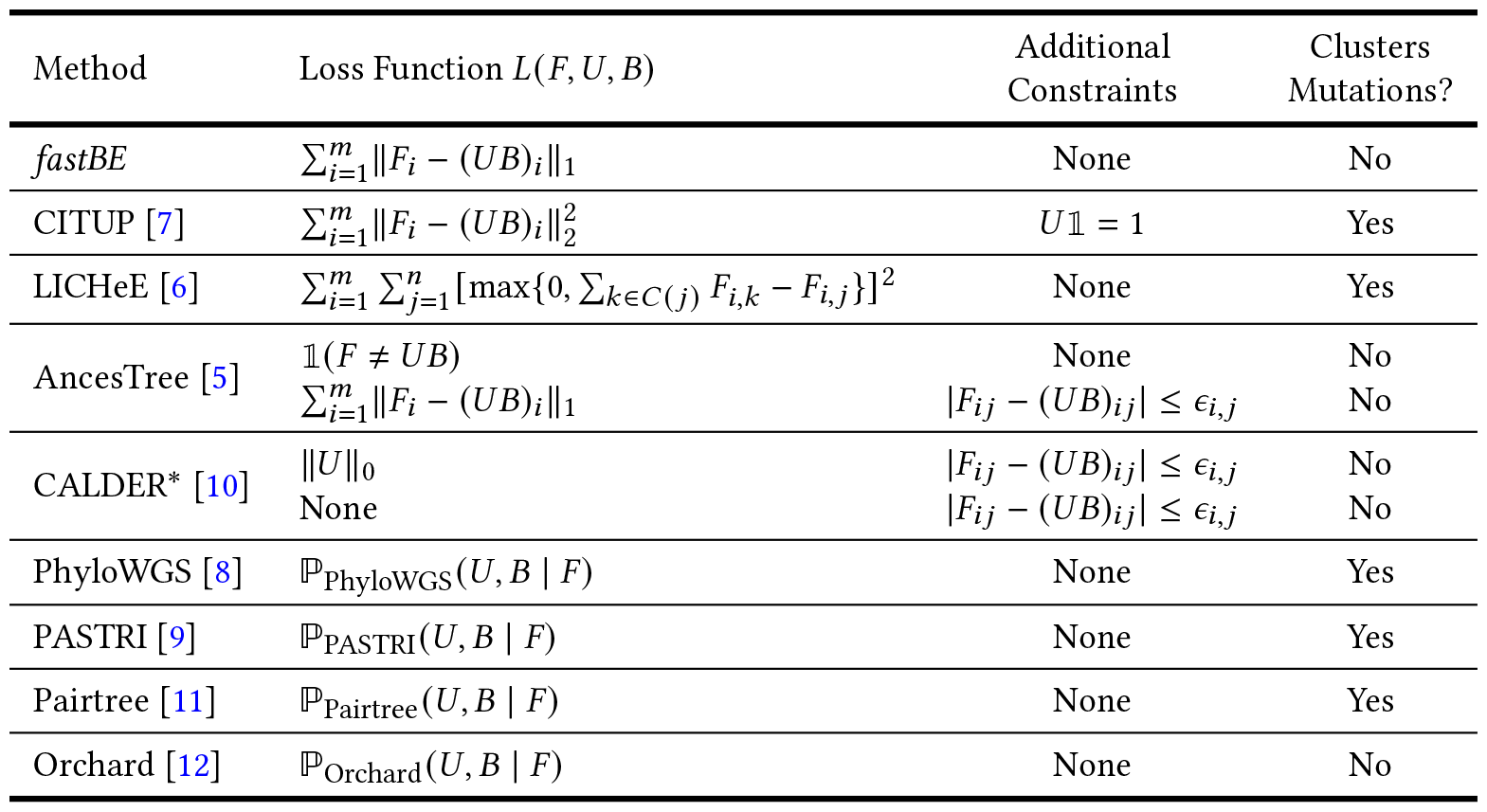
Summary of several variants of the variant allele frequency factorization problem studied in the literature. The loss functions *L*(*F, U, B*) and additional constraints beyond *U* ≥ 0, *U* 𝟙 ≤ 1 are noted for each method. Due to space constraints, we do not describe the regularization term(s) appearing in the loss function for several methods [6–9, 11], which penalize the total number of mutation clusters. ^*^CALDER enforces an additional hard constraint that the inferred matrices are *longitudinally consistent*, when provided with additional longitudinal information.

| Method | Loss Function $L(F, U, B)$ | Additional Constraints | Clusters Mutations? |
| --- | --- | --- | --- |
| *fastBE* | $\sum_{i=1}^m \ F_i - (UB)_i\ _1$ | None | No |
| CITUP [7] | $\sum_{i=1}^m \ F_i - (UB)_i\ _2^2$ | $U\mathbb{1} = 1$ | Yes |
| LICHeE [6] | $\sum_{i=1}^m \sum_{j=1}^n [\max\{0, \sum_{k \in C(j)} F_{i,k} - F_{i,j}\}]^2$ | None | Yes |
| AncesTree [5] | $\mathbb{1}(F \neq UB)$<br>$\sum_{i=1}^m \ F_i - (UB)_i\ _1$ | None<br>$\vert F_{ij} - (UB)_{ij}\vert \leq \epsilon_{i,j}$ | No<br>No |
| CALDER* [10] | $\ U\ _0$<br>None | $\vert F_{ij} - (UB)_{ij}\vert \leq \epsilon_{i,j}$<br>$\vert F_{ij} - (UB)_{ij}\vert \leq \epsilon_{i,j}$ | No<br>No |
| PhyloWGS [8] | $\mathbb{P}_{\text{PhyloWGS}}(U, B \mid F)$ | None | Yes |
| PASTRI [9] | $\mathbb{P}_{\text{PASTRI}}(U, B \mid F)$ | None | Yes |
| Pairtree [11] | $\mathbb{P}_{\text{Pairtree}}(U, B \mid F)$ | None | Yes |
| Orchard [12] | $\mathbb{P}_{\text{Orchard}}(U, B \mid F)$ | None | No |

The main focus of this work is the *structured regression* problem that appears when the clonal matrix *B* is fixed. Specifically, when *B* is fixed, the problem reduces to finding a usage matrix *U* such that the loss *L F, U, B* is minimized. We call this a *structured regression* problem as the aim is to *regress* the frequency matrix *F* against the clonal matrix *B* where *B* has a unique, *combinatorial structure* which we describe in Section 3.1.

### Problem 2

(The Variant Allele Frequency *L*-Regression Problem (*L*-VAFRP)). *Given a frequency matrix F and a clonal matrix B, find a usage matrix U such that the loss L*(*F, U, B*) *is minimized. We call this minimum loss L*^*^ (*F, B*).

In the special case where the loss 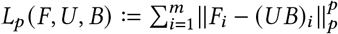 for *p* ∈ {1, 2}, this regression problem is solveable in polynomial time. More formally, since *u*_*i,j*_ ≥ 0 and 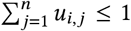 are linear constraints (1) on the matrix *U*, the *L*_1_ and *L*_2_ regression problems can be formulated as linear and convex quadratic programs respectively, which are both solveable in polynomial time. Throughout the remainder of this work, we will focus on the *L*_1_ regression and factorization problems, which we call the *variant allele frequency* ℓ_1_*-regression problem* (ℓ_1_-VAFRP) and *variant allele frequency* ℓ_1_*-factorization problem* (ℓ_1_-VAFFP), respectively. Further, though a slight abuse of notation, we will write 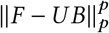 instead of *L*_*p*_ (*F, U, B*).

In contrast to the aforementioned approaches, which preprocess the read count data to obtain the observed frequency matrices *F*, probabilistic approaches such as PhyloWGS [8], PASTRI [9], Pairtree [11], and Orchard [12] explicitly model the read count data. Further, rather than minimize a loss, these methods attempt to sample from the posterior distribution of phylogenetic trees using a variety of sampling techniques. Since these probabilistic approaches are challenging to succinctly describe, we refer to the original publications [8, 9, 11, 12] for a complete description and denote their loss generically as ℙ(*U, B* | *F*) in Table 1.

General purpose linear programming software solves the ℓ_1_-VAFRP in polynomial time. However, linear programming solvers do not exploit the special structure of *B* and have worse asymptotic complexity as compared to the algorithm we derive in this work. Numerical methods from convex optimization such as the Alternating Direction Method of Multipliers and Projected Gradient Descent Method [31] can be used to solve the *L*_2_ variant allele frequency regression problem to a guaranteed optimality threshold. However, these blackbox methods are again quite slow and do not yield an exact solution. As such, [32] designed an improved algorithm that solves the *L*_2_ regression problem exactly in 𝒪 (*mn*^2^) time.

## 3 A structured regression model for the ℓ_1_-VAFFP

Structured regression and local search have played a pivotal role in the success of methods for performing distance based reconstruction of phylogenetic trees. In particular, state-of-the-art methods for distance based phylogenetics such as FastME [33] and FastTree [34, 35] work by locally exploring phylogenetic tree space (i.e. a tree space polytope [36, 37]) and regressing the observed distances against the tree to infer the unknown branch lengths, picking the tree which best explains the observed distances (Figure 1). Pivotal to the success of these methods are efficient algorithms [20, 38, 39] for computing and recomputing the solution to the structured regression problem. Importantly, these algorithms leverage the structured relationship between the branch lengths, the tree, and the induced distances.

**Figure 1:**
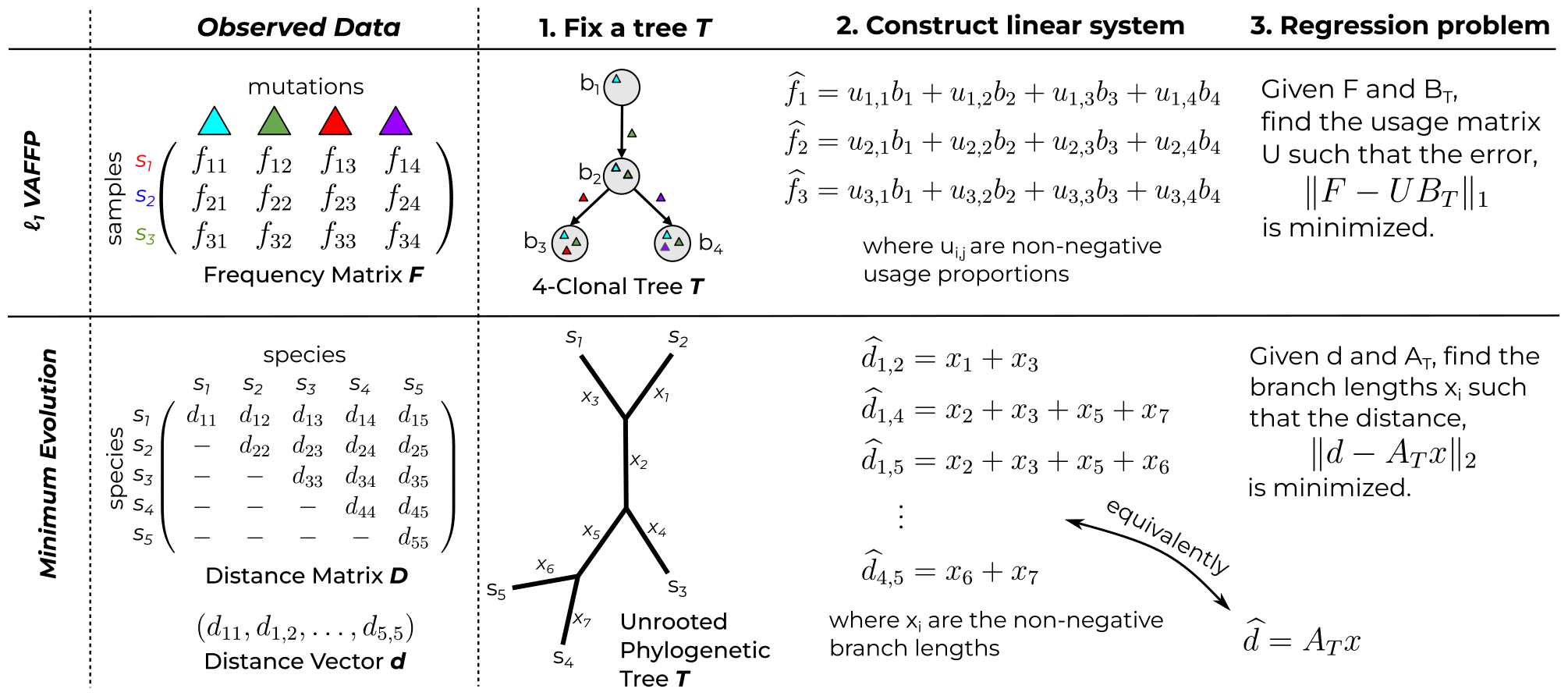
Analogous *structured regression* problems for a fixed tree topology *T* in the: **(top)** ℓ_1_-VAFFP framework for cancer evolution and the **(bottom)** minimum evolution framework for distance based phylogenetics. The usage matrix *U* describes the fraction of each clone across all samples, and the clonal matrix *B*_*T*_ describes the genotype of each clone in *T*. The matrix *A*_*T*_ describes the system of linear equations relating the branch lengths *x* to the distance vector *d*.

Despite the success of structured regression as a tool for distance based phylogenetics, such an approach has not yet been studied in the context of tumor evolution. Here, we derive an efficient algorithm for the ℓ_1_-VAFRP by exploiting the combinatorial structure of clonal matrices. In particular, we start by studying the structure of clonal matrices, both summarizing and extending existing results. Then, narrowing our focus, we derive an equivalent characterization of the ℓ_1_-VAFRP, which emphasizes the tree structure inherent in the problem. Finally, we use our characterization to design an efficient algorithm for the ℓ_1_-VAFRP which enables fast recomputation upon subtree prune-and-regraft (SPR) operations [23], providing the main theoretical result of our work, as stated below.

### Theorem 1.

*Given a clonal tree 𝒯 with n vertices and an m-by-n frequency matrix F, the minimum*

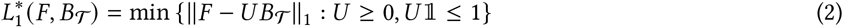

*can be found in 𝒪* (*mnd*) *time, where d is the depth of* 𝒯.

As mentioned above, our algorithm for solving the ℓ_1_-VAFRP is also able to efficiently recompute the minimum 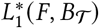 upon slight modifications to the tree topology, in the sense of the following corollary.

### Corollary 1.

*Given a clonal tree* 𝒯 *with n vertices and an m-by-n frequency matrix F, the following queries can be efficiently answered after 𝒪* (*mnd*) *pre-processing time using 𝒪* (*mnd*) *space*.

i. *For a subtree prune-and-regraft (SPR) operation on vertices i and j which results in a tree 𝒯* ^′^, *the minimum* 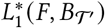 *can be queried in 𝒪* (*md* · max{*d* (*i*), *d* (*j*)}) *time*.
ii. *For the operation of attaching a new vertex j as a child of a vertex i to obtain a tree 𝒯*^′^ *and appending a corresponding column to the frequency matrix F to obtain F* ^′^, *the minimum* 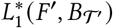 *can be queried in 𝒪* (*md* · *d* (*i*)) *time*.

Importantly, the depth *d* of a tree on *n* vertices is at most *n*− 1, implying our algorithm runs in quadratic (in *n*) time in the worst case and improves upon linear programming based approaches. However, for reasonable classes of trees, the complexity is much better. For example, the expected depth of a rooted spanning tree drawn uniformly at random from the complete graph *K*_*n*_ is of order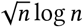– similar bounds can also be derived for the random spanning trees of an arbitrary graph *G* [40, 41]. All proofs can be found in Supplementary Proofs B.

### 3.1 Clonal trees and matrices

The most salient feature of clonal matrices is that they are (two-state) perfect phylogeny matrices [5], and this allows us to tap into the theory which studies such matrices. However, the class of clonal matrices is much more restrictive: there are a handful of useful results concerning clonal matrices which are not applicable to perfect phylogeny matrices. For example, a perfect phylogeny matrix need not be square and thus it is not necessarily invertible, while clonal matrices are always invertible [5]. Due to space constraints, we move the discussion of clonal matrices and trees to Supplementary Results A.1.

We conclude this section with an algebraic description of the inverse of *B*, which was previously described in [32], and will serve as a key technical ingredient for deriving an equivalent formulation of the ℓ_1_-VAFRP. Define the adjacency matrix *A* of 𝒯 as the *n*-by-*n* matrix such that *A*_*i,j*_ = 1 if *i* is a parent of *j* in 𝒯 and 0 otherwise. Then, the inverse of *B* is (*I* − *A*), where *I* is the identity matrix.

#### Lemma 1.

*For any clonal matrix B associated with a clonal tree 𝒯 having adjacency matrix A, B* = (*I* − *A*)^−1^. *Consequently*, [*B*^−1^*v*]_*i*_ = *v*_*i*_ − *v*_*δ* (*i*)_ *if i* ≠ *r* (𝒯), *and* [*B*^−1^*v*]_*i*_ = *v*_*i*_ *when i* = *r* (𝒯).

### 3.2 An equivalent formulation of the VAF ℓ_1_-regression problem

In this section, we show that the ℓ_1_-VAFRP is equivalent to a constrained vertex labeling problem on the clonal tree 𝒯 associated with *B*. Specifically, we prove that the ℓ_1_-VAFRP is equivalent to a special case of the *dot product tree labeling problem*, defined as follows, for the case when *m* = 1 and *F* = *f* ^*T*^. The general case of *m >* 1 follows from the separability of the objective 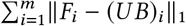.

#### Problem 3

(Dot Product Tree Labeling Problem (DPTLP)). *Given a rooted tree 𝒯 with vertices* [*n*] *and a vector w* ∈ ℝ^*n*^, *find a non-negative vector x* ∈ ℝ^*n*^ *such that x*^*T*^ *w is maximized and* |*x*_*i*_ − *x*_*δ* (*i*)_ | ≤ 1 *for all i* ≠ *r* (𝒯), *where δ* (*i*) *is the parent of vertex i in 𝒯*.

To derive the equivalence between the ℓ_1_-VAFRP and the DPTLP, we start by writing the ℓ_1_-VAFRP as a linear program (LP) in standard form [31]. Using the usual trick [31] for converting the ℓ_1_ norm to a linear objective with linear constraints, we write the ℓ_1_-VAFRP as a LP (Figure 2). Then, we write out the dual problem by associating a dual variable *α*_*i*_ with the constraint in (3), a dual variable *β*_*i*_ with the constraint in (4), and a dual variable *γ* with the constraint in (5). Thus obtaining our dual LP (Figure 2).

**Figure 2:**
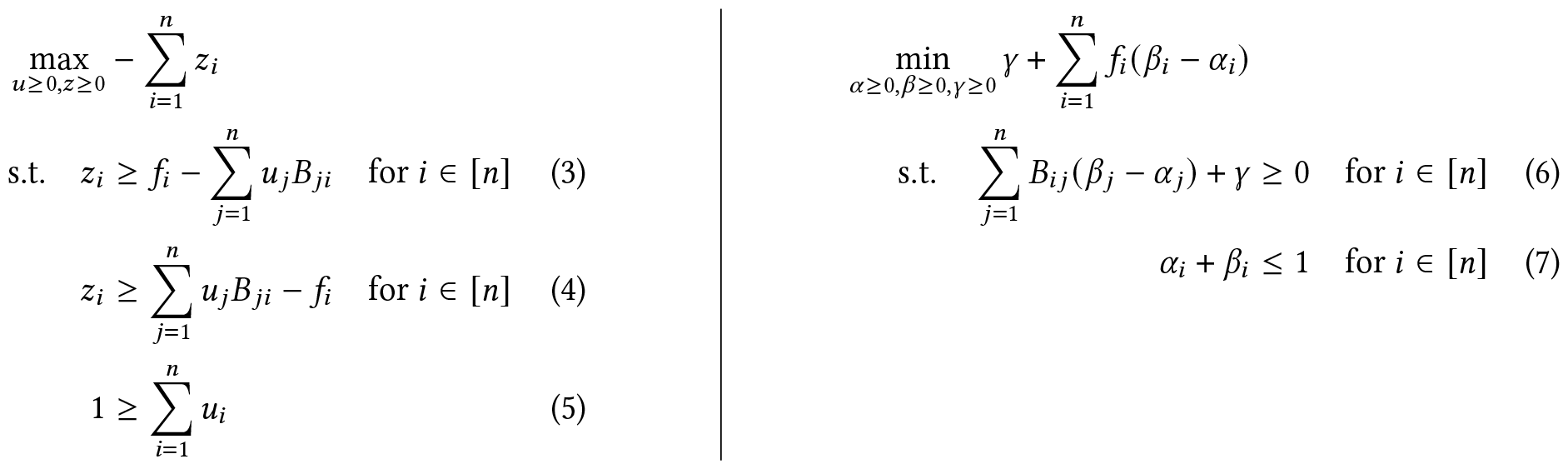
**(Left)** The primal LP and **(right)** the dual LP for the ℓ_1_-VAFRP

To simplify the dual form of the LP, we perform a change of variables by setting *λ*_*i*_ = *β*_*i*_ − *α*_*i*_. Since *α*_*i*_ and *β*_*i*_ are non-negative and their sum is bounded by 1, *λ*_*i*_ ∈ [− 1, 1]. Then, writing the constraints in matrix form and using a slack variable *ψ* to remove the inequality constraint, we have the following equivalent, dual LP.

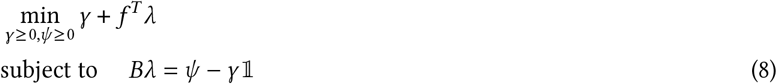

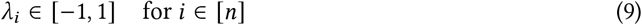

Applying Lemma 1 to the matrix *B* in (8) and invoking LP duality then proves the following theorem.

#### Theorem 2.

*Given a length n frequency vector f and an n-by-n clonal matrix B corresponding to a clonal tree with root vertex* 1, *the minimum of* ∥ *f* ^*T*^ − *u*^*T*^ *B*∥_1_ *over all usage vectors u* ∈ ℝ^*n*^ *is equal to the maximum of*

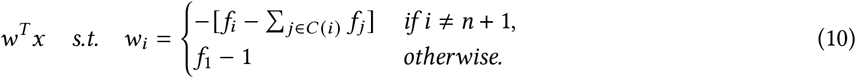

*over all non-negative vectors x* ∈ ℝ^*n*+1^ *such that* |*x*_1_ − *x*_*n*+1_| ≤ 1 *and* |*x*_*i*_ − *x*_*δ* (*i*)_ | ≤ 1 *for all i* ∈ [*n*].

In other words, the above theorem states that the ℓ_1_-VAFRP is equivalent to a special case of the DPTLP where we append a parent labeled *n* +1 to the root vertex and appropriately set the vector *w* ∈ ℝ^*n*+1^. Interestingly, the *sum condition* [5–7, 24], that is, the requirement that

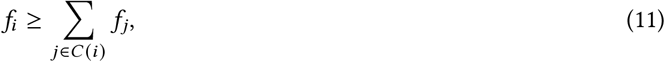

appears almost unexpectedly in (10). Using the appearance of the sum condition in (10), we extend the theory of El-Kebir et al. [5] which states that we can find a usage vector *u*^*T*^ such that *f* ^*T*^ = *u*^*T*^ *B* if and only if the sum condition (11) is satisfied. In particular, we show that the total violation of the sum condition also provides a lower bound on the ℓ_1_ error.

#### Corollary 2.

*Let f be a frequency vector of length n, let B be an n-by-n clonal matrix, and let* 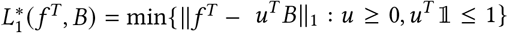. *then* 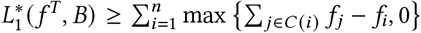, *or the total violation of the sum condition* (11). *Thus*, 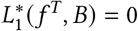 *if and only if the sum condition is satisfied*.

Finally, we observe that as a consequence of the above corollary, the sum condition is somewhat redundant when minimizing the ℓ_1_ error ‖*F*− *UB* ‖_1_. This is because by the above corollary, the ℓ_1_ error is an upper bound on the total violation on the sum condition, implying that minimizing the ℓ_1_ error forces the total violation of the sum condition to zero.

### 3.3 An algorithm for the DPTLP

In this section, we develop an efficient algorithm for the DPTLP. There are two key ideas underlying our algorithm. The first idea is that exploiting the tree structure enables us to express the solution for the subtree rooted at a vertex *i* in terms of the solution for the subtrees rooted at the children of *i*. This expression is derived using standard techniques for dynamic programming on trees [42]. The second idea is more technical, and is based on the observation that the solution of this recurrence is a concave piecewise linear function, which we represent compactly as a list of size 𝒯 (*d*), where *d* is the depth of 𝒪. Combining these two ideas yield an O(*nd*) algorithm for the DPTLP, as stated below.

#### Theorem 3.

*Given a rooted tree 𝒯 of depth d with vertices* [*n*] *and a vector w* ∈ ℝ^*n*^, *the DPTLP can be solved in 𝒪* (*nd*) *time with the following recurrence*,

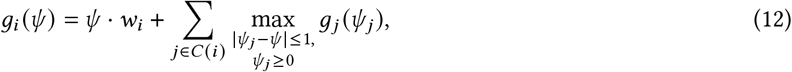

*where g*_*i*_ (*ψ*) *is the optimal solution to the DPTLP for the subtree rooted at vertex i, when vertex i is assigned the label ψ*. We will prove this theorem by first deriving a recurrence relation for the solution of the DPTLP. Assume *g*_*i*_ (*ψ*) is the optimal solution to the DPTLP for the subtree rooted at *i* when the root vertex *i* is assigned the label *ψ*. Then, *g*_*i*_ (*ψ*) satisfies the following recurrence relation:

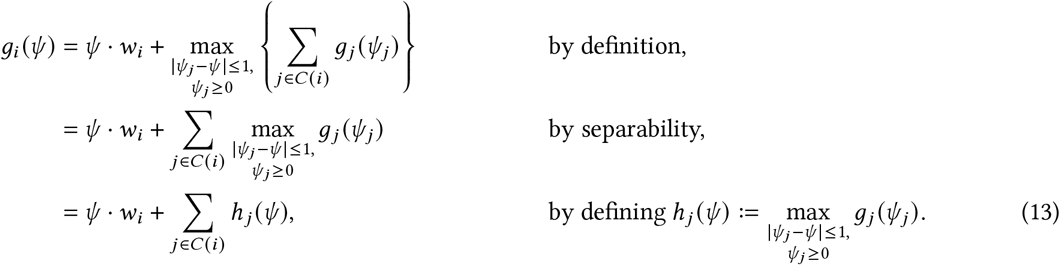

Importantly, to solve the DPTLP, it is necessary and sufficient to compute max_*ψ*_ ≥_0_ *g*_*r*_ *ψ* for the root vertex *r* = *r*(𝒯). Unfortunately, however, the straightforward technique [42] of storing a dynamic programming table for this recurrence will not work because the number of possible values of *ψ* ≥ 0 is infinite. Thus, we need to find an alternative way to describe and represent the functions *g*_*j*_ and *h* _*j*_ appearing in this recurrence.

The key mathematical idea underpinning our algorithm is that the functions *g*_*j*_ and *h* _*j*_ appearing in the recurrence are not arbitrary. Rather, they form a special class of functions which admit a compact representation and are convenient to work with. In particular, the functions *g*_*j*_ and *h* _*j*_ are concave (and thus continuous) piecewise linear functions with a finite number of breakpoints at coordinates {1, 2, …, *k*}, formally defined below.

#### Definition 3.

*Let ℒ*_*k*_ *be the set of concave piecewise linear functions with breakpoints at integers* 1, …, *k; i*.*e*., *a continuous function f*∈ℒ _*k*_ *if f is linear with slopes s*_1_ ≥ *s*_2_ ≥… ≥ *s*_*k*+1_ *on the intervals I*_1_ = [0, 1), *I*_2_ = [1, 2), …, *I*_*k*+1_ = [*k*, ∞) *respectively*.

Note that ℒ_*k*_ form a strictly increasing nested sequence ℒ_0_ ⊂ℒ_1_⊂ … of sets of piecewise linear functions. Further, every function *f* in the class ℒ_*k*_ is represented by a tuple (y, *s*_1_, …, *s*_*k* +1)_ of size *k*+ 2 giving the intercept y and slopes *s*_*i*_ of each piece of *f*.

Now, we will prove that *g*_*j*_ and *h* _*j*_ are in the class ℒ_*d*_, where *d* is the depth of 𝒯. To start, notice that if *i* is a leaf vertex, then *g*_*i*_ (*ψ*) = *ψ* · *w*_*i*_ is a linear function and is thus in ℒ_0_. This provides us with the necessary base case required to prove that *g*_*j*_ is in ℒ_*d*_. Next, we prove the inductive step of our claim: if *g*_*j*_ ∈ ℒ_*k*_, then *h* _*j*_ ∈ ℒ_*k*+1_. The results then follow by induction on the depth of 𝒯. We start with the following description of *h* _*j*_ in terms of *g*_*j*_.

#### Lemma 2.

*Suppose f* ∈ *L*_*k*_ *and is represented by the tuple* (y, *s*_1_, *s*_2_, …, *s*_*k*+1_). *Let i*^*^ *be the largest index i such that s*_*i*_ ≥ 0. *If s*_*i*_ *<* 0 *for all i, we set i*^*^ = −∞ *and if s*_*i*_ ≥ 0 *for all i, we set i*^*^ = ∞. *Then*,

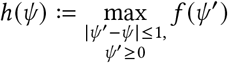

*is in* ℒ_*k*+1_ *and satisfies*

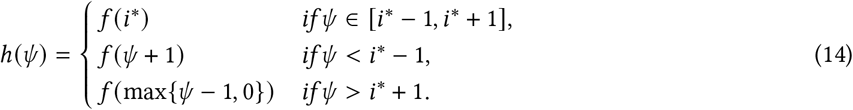

*Proof*. If *i*^*^ = ∞, all slopes *s*_*i*_ are non-negative and the function *g*(*ψ* ^′^) is non-decreasing. Thus, the maximum of *g*(*ψ* ^′^) over any interval [*ψ* − 1, *ψ* + 1], is achieved at the interval’s right most value *ψ* + 1.

If *i*^*^ = − ∞, all slopes *s*_*i*_ are negative and the function *g* (*ψ* ^′^) is strictly decreasing. Thus, the maximum of *g* (*ψ* ^′^)over any interval [*ψ* − 1, *ψ* +1], is achieved at the interval’s left most value *ψ* − 1. However, since *ψ* ^′^ is constrained to be non-negative, if *ψ <* 1 the maximum is achieved at *ψ* ^′^ = 0.

If *i*^*^ ≠∞, − ∞, then *g* (*ψ* ^′^) is non-decreasing on the interval [0, *i*^*^] and non-increasing on the interval [*i*^*^,∞). Further, the maximum of *g* (*ψ* ^′^) over all non-negative *ψ* ^′^ is bounded and equal to *g* (*i*^*^). The result then follows by a case analysis on the value of *ψ*. If *ψ* is in [*i*^*^ −1, *i*^*+^ 1], then we can take *ψ* ^′^ = *i*^*^ and achieve the maximum. If *ψ < i*^*^ − 1, then the function *g* (*ψ* ^′^) is non-decreasing on the interval [*ψ* − 1, *ψ* +1] and the maximum is achieved at the interval’s right most value *ψ* + 1. Symetrically, if *ψ > i*^*^ + 1, then the function *g (ψ*′) is non-increasing on the interval [*ψ* − 1, *ψ* + 1] and the maximum is achieved at the interval’s left most value *ψ* − 1, which is always non-negative since *ψ >* 1. As this covers all possible cases, this proves that *h* has the form (14).

To see that *h*(*ψ*) is continuous, observe that

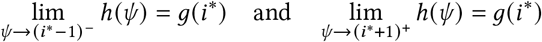

at the only candidates for discontinuity, *i*^*^ − 1 and *i*^*^ + 1. To see that it is concave, note that

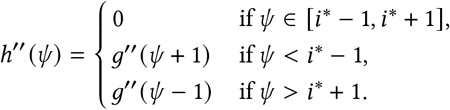

As *g ψ* is concave, its second derivative *g*^′′^ *ψ* is non-positive, which implies that *h ψ* is also concave. Since *h* is trivially piecewise linear by (14), the proof is complete.

Stated in terms of the tuple representations of *h* _*j*_ and *g*_*j*_ as tuples, we have the following equivalent result.

#### Proposition 1.

*Suppose f* ∈ *L*_*k*_ *and is represented by the tuple* (y, *s*_1_, *s*_2_, …, *s*_*k*+1_). *Let i*^*^ *be the largest index i such that s*_*i*_ ≥ 0. *If s*_*i*_ *<* 0 *for all i, we set i*^*^ = −∞ *and if s*_*i*_ ≥ 0 *for all i, we set i*^*^ = ∞. *Then*,

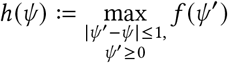

*is in* ℒ_*k*+1_ *and is represented by the tuple*

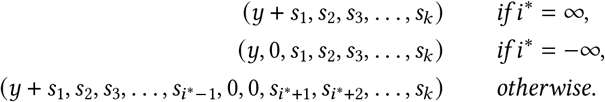

In summary, we have shown that *i)* if *i* is a leaf, *g*_*i*_ is in ℒ_0_, and *ii)* if *g*_*j*_ is in ℒ_*k*_, then *h* _*j*_ is in ℒ_*k*+1_. The final step is to show that if the function *h* _*j*_ is in ℒ_*k*_ for all children *j* of *i*, then *g*_*i*_ is also in ℒ_*k*_. This follows from the observation that the class ℒ_*k*_ is closed under addition. The result is summarized below.

#### Proposition 2.

*Let g*_*i*_ (*ψ*) *be the optimal solution of the DPTLP for the subtree* T_*i*_ *rooted at vertex i such that i is assigned the label ψ*. *Then, g*_*i*_ *is in* ℒ_*d*_.

We are now ready to prove the main result of this section.

*Proof of Theorem 3*. The result follows by induction on *n*, the number of vertices in 𝒯. In particular, assume that the representation of the function *g*_*i*_ for the root vertex *i* is computable in 𝒪 (*nd*) time for all trees with fewer than *n* vertices and depth at most *d*. Clearly, this holds for *n* = 1.

Then, let *r* be the root of 𝒯 and let *C* (*r*) be the set of children of *r*. Let 𝒯 _*j*_ denote the subtree rooted at *j* ∈ *C* (*r*) and *n* _*j*_ denote the size of 𝒪 _*j*_. By the inductive hypothesis, we can compute the representation of *g*_*j*_ for *j* ∈ *C* (*r*) in 𝒪 (*dn* _*j*_) time, since the depth of the subtree rooted at *j* ∈ *C* (*r*) is at most *d* + 2. Using Proposition 1, we can compute the representation of *h* _*j*_ for all *j* ∈ *C* (*r*) in 𝒪 (*d* Σ_*j* ∈*C* (*r*)_ *n* _*j*_) = 𝒪 (*nd*) time as these functions are represented by tuples of length at most *d* + 3.

Finally, we compute the representation of *g*_*r*_ by observing that ℒ_*d*_ is closed under addition and that the representation of *g*_*r*_ is easily computed from the representations of *h* _*j*_. In particular, summing the tuples representing *h* _*j*_ coordinate-wise, we obtain the representation of *g*_*r*_.

Then, by reducing the ℓ_1_-VAFRP to the DPTLP using Theorem 2 and applying Theorem 3, we complete the proof of the first part of Theorem 1 for the special case where *m* = 1. The general case of *m >* 1 follows by noting that the objective 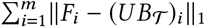 is separable and that each term ∥*F*_*i*_ − (*UB*_𝒯_)_*i*_ ∥_1_ can be minimized independently.

To see Corollary 1, notice that after pruning and regrafting any subtree 𝒯_*i*_ as a child of a vertex *j, g*_*k*_ only changes if the vertex *k* is on the path from root *r*(𝒯) to the vertex *δ* (*i*) or the vertex *j* – this follows from equation (13). Since there are at most 2 max {*d* (*j*), *d* (*i*) vertices on these paths, this observation yields the statement (i) in Corollary 1, as long as the representations of *g*_*k*_ are stored. The second statement (ii) in Corollary 1 follows by a similar argument.

## 4 A deterministic search algorithm for the 𝓁_1_-VAFFP

Here we describe a deterministic search algorithm, *fastBE*, for the 𝓁_1_-VAFFP which builds upon the fast regression algorithm described in Section 3 and is inspired by the beam search techniques used by Orchard [12]. In particular, our algorithm iteratively constructs the inferred tree one mutation at a time, choosing the best vertex placement for each mutation in the current tree, while allowing added vertices to “adopt” children from their parent. This procedure is described formally as follows:

1. Fix an order *O* = {*o*_1_, …, *o*_*n*_} in which to append the mutations *O* and initialize the starting tree as 𝒯_1_ =({ *o*_1_}, {}) with root *r*(𝒯) _1_ = *o*_1_.
2. For *i* = 2, …, *n*:
  a. Let the current tree 𝒯= 𝒯_*i*−1_ and the current frequency matrix *F*^′^ be the submatrix of *F* spanned by the columns *o*_1_, …, *o*_*i*_.
  b. Find the parent *p* ∈ *V* (𝒯) and the subset of children *S* ⊆ δ _𝒯_ (*p*) such that the tree 𝒯 ′ obtained from attaching *o*_*i*_ as a child of *p* in 𝒯 and adopting the subtrees rooted at the vertices in *S* as children of *o*_*i*_ minimizes the loss *L*^*^ (*F* ^′^, *B*_𝒯_′).
  c. Set the next tree 𝒯_*i*_ = 𝒯 ′.
3. Output the final tree 𝒯_*n*_.

Since at every iteration *i* of the algorithm there are a total of 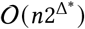 where 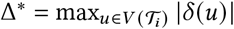 placements of mutation *o*_*i*_ to consider, the running time of this algorithm is dominated by the time to compute the objective function 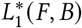 over all such placements. Using the efficient recomputation procedure outlined in the latter part of Theorem 1, we can avoid the naïve approach which takes 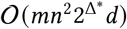 time and instead perform all such computations in 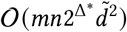 time, where 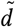 is the average depth of the tree 𝒯_*i*_. Details on these steps are supplied in Supplementary Section C.4.

## 5 Results

### 5.1 Runtime comparison to linear programming solvers

We compared an implementation of our structured regression algorithm for the 𝓁_1_-VAFRP to a linear programming (LP) approach on simulated data. In particular, we implemented the natural primal LP formulation solving the 𝓁_1_-VAFRP (Section 3.2) using two commercial LP solvers: Gurobi v9.0.3 [43] and CPLEX v22.1.0 [44]. We generated 264 pairs of frequency matrices *F* and clonal matrices *B* as described in Supplementary Section C.1, and measured the wall-clock runtime of our algorithm and the LP solvers on these simulated instances. Excluding the time required to construct the LP – which would unfairly penalize the LP solvers – we found that our algorithm was a mean of 95.6 times faster than Gurobi and 105.1 times faster than CPLEX (Supplementary Figures 1, 2).

Next, we tested the warm start capability of our structured regression algorithm for the 𝓁_1_-VAFRP upon perturbations to the topology of the input tree. For each of the 264 instances constructed above, we measured the time to solve the 𝓁_1_-VAFRP for 25,000 trees obtained by applying a single random SPR operation to the input clonal tree. We performed this measurement both in the setting where we employ our regression algorithm as a black-box (the *cold start* setting) and the setting where we used the recomputation procedure outlined in Corollary 1 (the *warm start* setting). We found that our algorithm was a mean of 6.2 times faster in the warm start as opposed to cold start settings (Supplementary Figures 3, 4). This implies that our regression algorithm possesses another advantage over naïve LP approaches, which do not provide any warm starting capabilities.

### 5.2 Evaluation of *fastBE* on simulated data

We evaluated our factorization algorithm, *fastBE*, on simulated data, and compared it to four other state-of-the-art factorization algorithms: Pairtree [11], Orchard [12], CALDER [10], and CITUP [7]. To perform our evaluation, we simulated ground truth clone trees and usage matrices, and measured the ability of each algorithm to reconstruct this ground truth. To construct each simulated instance, we generated a clone tree 𝒯 and usage matrix *U*, computed the frequency matrix *F* = *UB*, and sampled both variant and non-variant reads from *F* at 40× coverage. Complete details describing the simulations, parameters, and evaluation metrics are provided in Supplementary Sections C.1, C.2, C.3.

On simulated instances with few clones (*n* = 3, 5, 10) and samples (*m* = 5, 10, 25), all algorithms terminated in under 24 hours on the majority of the 108 simulated instances (*fastBE*: 108 /108, Pairtree: 108 /108, Orchard: 108 /108, CALDER: 106 /108, CITUP: 107/ 108). *fastBE*, Pairtree, and Orchard accurately recovered pairwise relationships in this setting (Figure 3; Supplementary Figure 5), with *fastBE*, Pairtree, and Orchard performing nearly identically for the *n* = 10 clone setting in terms of mean F1-score (*fastBE*: 0.965, Pairtree: 0.972, Orchard: 0.965). In contrast, CITUP and CALDER struggled to accurately reconstruct pairwise relationships (Figure 3; Supplementary Figure 5) when there were 5 or more clones. In terms of recovering the ground truth usage matrix *U* and frequency matrix *F, fastBE* significantly outperformed CITUP and CALDER, while performing similarly to Pairtree and Orchard (Supplementary Figure 6).

**Figure 3:**
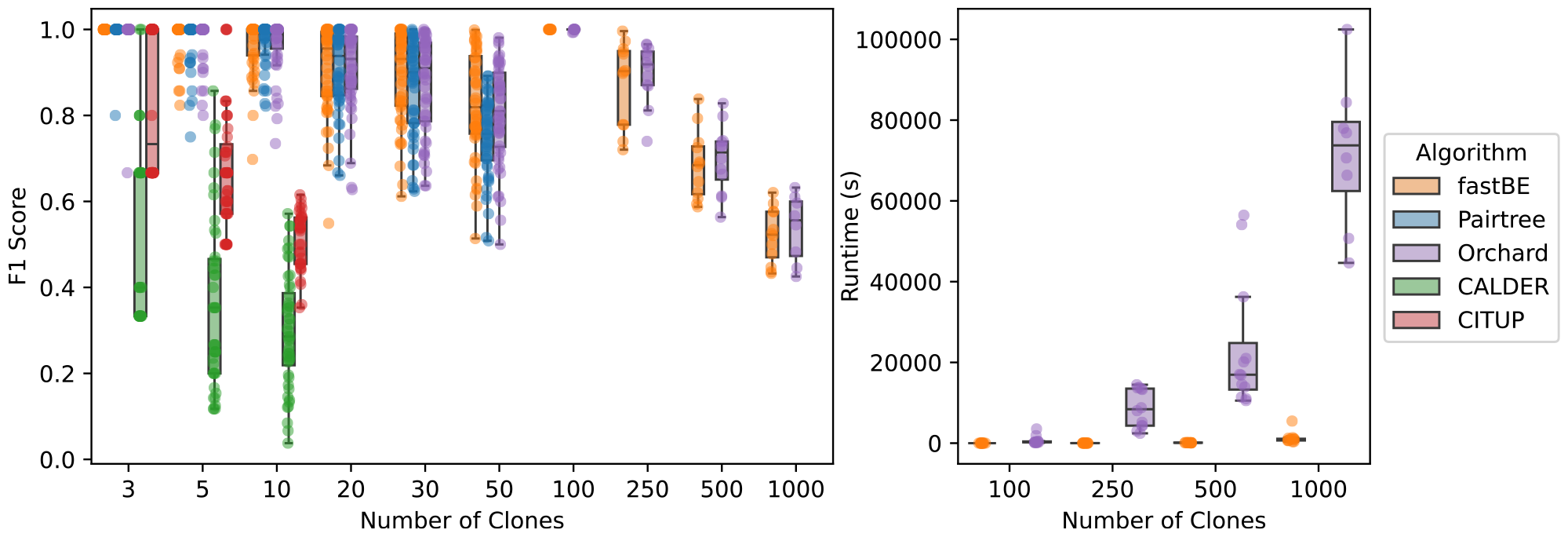
The accuracy of reconstructing pairwise relationships for the phylogenetic trees inferred by *fastBE*, Pairtree [11], Orchard [12], CALDER [10], and CITUP [7] on simulated data. **(Left)** The F1-score versus the number of clones. **(Right)** The wall-clock runtime on instances with ≥100 clones versus the number of samples. Methods that did not scale to instances with many clones are excluded from the plot.

On simulated instances with a modest number of clones (*n* = 20, 30, 50) and samples (*m* = 25, 50), CITUP and CALDER were unable to terminate on the majority of the simulated instances within 24 hours – consistent with the findings of [11, 12] – and were excluded from our evaluation. On instances with *n* = 20, 30 clones, *fastBE*, Pairtree, and Orchard performed similarly in terms of reconstructing pairwise relationships (Figure 3; Supplementary Figure 7). However, on instances with *n* = 50 clones, *fastBE* and Orchard outperformed Pairtree (mean F1 *fastBE*: 0.825, Pairtree: 0.749, Orchard: 0.805) in reconstructing pairwise relationships (Supplementary Figure 13). In terms of recovering the ground truth usage matrix *U* and frequency matrix *F*, all methods performed quite well in recovering *F*, but Pairtree was less accurate in recovering *U* and *F* when the number of clones was large (Supplementary Figure 8). Finally, *fastBE* was an order of magnitude faster than Pairtree and Orchard, running for an average of 1.21 seconds on instances with *n* = 50 clones (Supplementary Figure 13).

In the regime with a large number of clones (*n* = 100, 250, 500, 1000) and samples (*m* = 50, 100), only *fastBE* and Orchard were able to terminate within a 24 hour time limit. On these instances, *fastBE* and Orchard perform nearly identically in terms of recovering ground truth pairwise relationships (mean F1 *fastBE*: 0.773, Orchard: 0.783). The methods also had similar performance in recovering the ground truth usage matrix *U* and frequency matrix *F* (Supplementary Figure 8). In terms of runtime, however, *fastBE* was several orders of magnitude faster than Orchard (Figure 3; Supplementary Figure 13). For example, *fastBE* took a mean of 1229.8 seconds to run on instances with *n* = 1000 clones and terminated on all such instances, whereas Orchard took a mean of 71749.1 seconds and terminated on 19/24 such instances when allotted 48 hours and a dedicated 32-core processor.

We also found that *fastBE* was more *sample efficient* than other methods, requiring fewer samples to recover the ground truth clonal relationships. In particular, the pairwise reconstruction accuracy strictly improved for *fastBE*, Pairtree, and Orchard as the number of samples increased (Figure 3; Supplementary Figure 12), while this was not necessarily the case for CALDER and CITUP (Supplementary Figures 9, 10). However, the reconstruction accuracy improved for *fastBE* more quickly than Pairtree and Orchard (Figure 3; Supplementary Figure 12) as the number of samples increased, and *fastBE* obtained near perfect recovery (median F1: 0.987) with the number of samples *m* ≥ 50 and the number of clones *n <* 100. This observation led us to investigate the reconstruction accuracy as the ratio of samples to clones increased. Interestingly, we observed a sharp transition in pairwise reconstruction accuracy for *fastBE* as the ratio of samples to clones approached one (Supplementary Figure 11).

### 5.3 Analysis of B progenitor acute lymphoblastic leukemia patient samples

We applied *fastBE* to infer phylogenetic trees from multi-sample bulk DNA sequencing data of fourteen patients with B progenitor acute lymphoblastic leukemia (B-ALL) [16]. This dataset was sequenced at greater than 200× coverage using targeted sequencing, containing a median of 42 (min: 13, max 90) samples per patient. Originally, mutation clusters and phylogenetic trees were inferred from this dataset using Pairtree [11]. To perform a fair comparison of *fastBE* against Pairtree, we applied *fastBE* to the (median: 8, min: 3, max: 17) mutation clusters inferred by Pairtree for each patient, obtaining phylogenetic trees on the exact same mutation clusters. We found that on all of the 14 of the patients, both *fastBE* and Pairtree inferred distinct trees, having a median of at least 8 distinct edges (Supplementary Figures 14). For both methods, the normalized frequency matrix estimation error 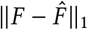was less than 1% in all but one case (Supplementary Figures 15), though the trees inferred by *fastBE* had modestly lower frequency matrix estimation error.

We quantified the differences between the phylogenetic trees inferred by *fastBE* and Pairtree by examining violations of the *sum condition* (equation (11)) [5–7, 24] in the phylogenetic trees. Specifically, the *sum condition* requires that the frequency *F*_*i,j*_ of a mutation *j* gained at a clone in sample *i* is greater than or equal to the sum ∑_*k* ∈*C* (*j*)_ *F*_*i,k*_ of the frequencies of the mutations gained at the clone’s children. The sum condition follows from the perfect phylogeny assumption, which states that each mutation is gained at most once and never lost. Consequently if a mutation is present in a clone, then the mutation is present in all the clone’s children. Importantly, all of the methods benchmarked [7, 10–12] make the perfect phylogeny assumption, and thus if the frequency matrix is correctly measured, the inferred trees should satisfy the sum condition. For a sample *i* and mutation *j*, we define the *violation V*_*i,j*_ = max {∑_*k* ∈*C*(*j*)_ *F*_*i,k*_ − *F*_*i j*_, 0} of the sum condition. The *total violation V* of the sum condition is the sum *V* =∑_*i,j*_ *V*_*i,j*_ of the violations over all samples and mutations. We found that the phylogenies inferred by *fastBE* had a lower total violation *V* (mean *V* of 1.44 over 14 patients) compared to Pairtree (mean of *V* of 2.40) (Figure 4a) across all 14 patients.

**Figure 4:**
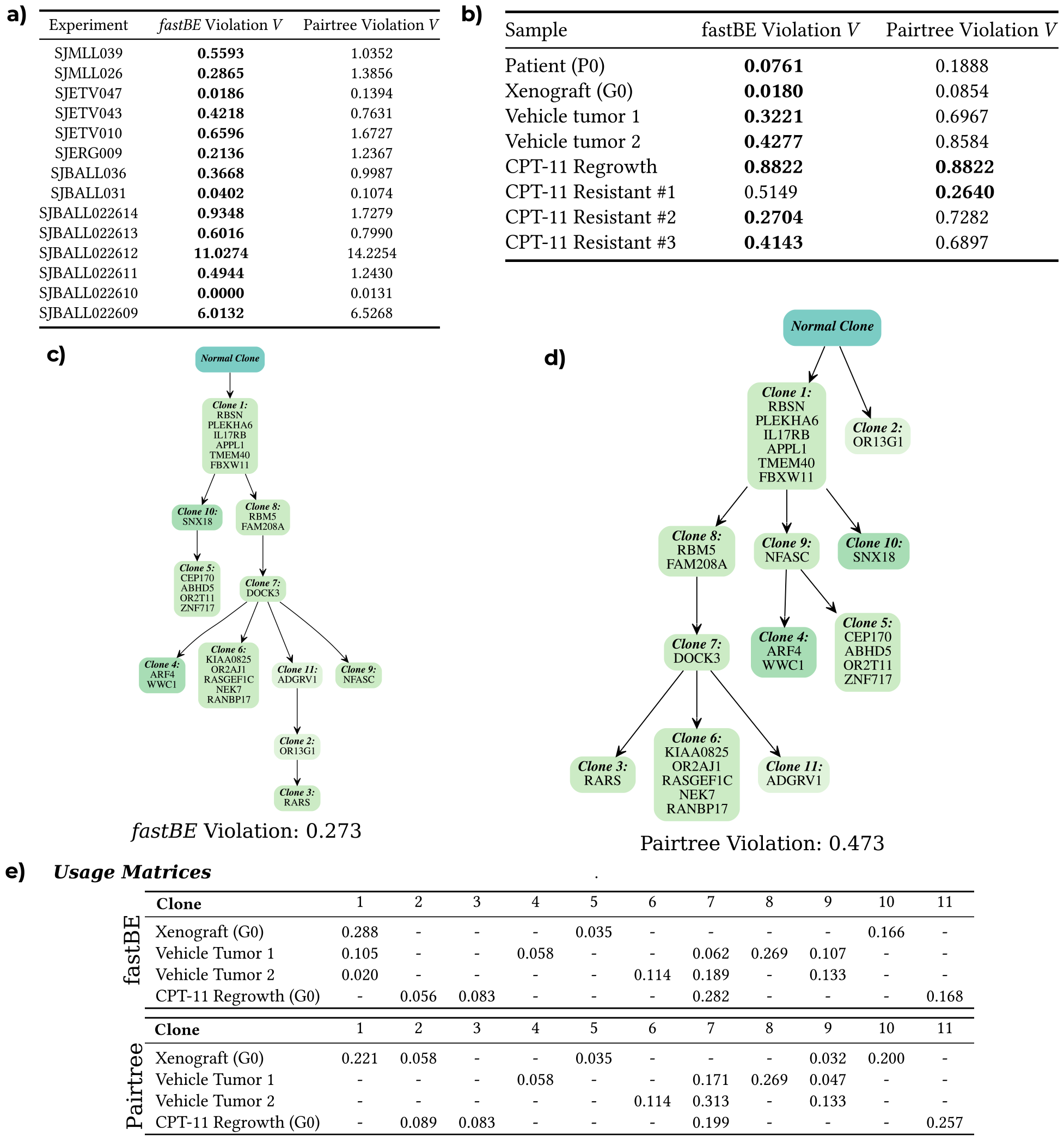
**a)** The total violation 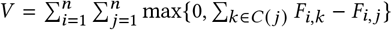 of the sum condition for the phylogenies inferred by *fastBE* and Pairtree [11] on data from fourteen patients with B progenitor acute lymphoblastic leukemia. **b)** The per-sample violation 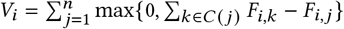 of the sum condition for the phylogenies inferred by *fastBE* and Pairtree from the multi-sample bulk DNA sequencing data of the POP66 colorectal cancer model [45]. The phylogenies inferred by **c)** *fastBE* and **d)** Pairtree for the CSC28 colorectal cancer model. Each vertex corresponds to a distinct clone and is labeled by the set of mutations which occur on the edge directed into the clone. **e)** Usage matrices *U* inferred from the phylogenetic trees output by *fastBE* and Pairtree. Entries marked as ‘−’ correspond to usage values of 0.0.

### 5.4 Analysis of patient-derived colorectal cancer models

Finally, we compared *fastBE* and Pairtree on two patient-derived xenograft models of colorectal cancer, POP66 and CSC28 from [45], from which multiple bulk samples underwent whole-exome sequencing. The POP66 model contained eight samples collected in the parent tumor (P0), first generation xenograft (G0), and regrowth xenografts, and 25 clones were inferred across these samples in [45]. The CSC28 model consisted of four samples collected in the first generation (G0) and regrowth xenografts, and contained 11 clones were inferred in these samples in [45]. When applying *fastBE* to the CSC28 and POP66 models, we fixed the root clone to contain all mutations with VAF near 0.5 across all samples.

We found that the phylogenetic trees inferred by *fastBE* were quite different from those inferred by Pairtree in terms of both their implied pairwise relationships and overall structure (Figure 4). For example, while both CSC28 phylogenies had the same clone 1 as a child of the normal clone, the mutation ORG13G was a child of the normal clone in the Pairtree phylogeny, implying a polyclonal tumor origin. Further, while clone 4 is a child of clone 7 in the *fastBE* phylogeny, clone 4 is on a separate branch from clone 7 in the Pairtree phylogeny. A similar story appears for the POP66 phylogenies, where the differences are even further exaggerated due to the large number of clones.

Next, we quantified violations of the sum condition in the phylogenetic trees inferred by *fastBE* and Pairtree. We found that *fastBE* had substantially lower total violation than Pairtree on both the CSC28 (*fastBE*: 0.273, Pairtree: 0.473) and POP66 (*fastBE*: 3.280, Pairtree: 4.770) phylogenies. This trend held separately across all samples (Figure 4b), and was most pronounced for the “CPT-11 Resistant” samples [45] in POP66.

Finally, we compared the inferred clonal proportions in each sample, a measure of the heterogeneity within the tumor samples. We define the heterogeneity 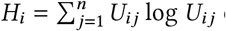 of a sample *i* as the Shannon diversity index, a commonly used measure of species diversity in the ecology literature [46]. (Note that 0 log 0 = 0.) For CSC28, we found similar sample heterogeneity estimates for both *fastBE* and Pairtree. For POP66, we found similar heterogeneity estimates for most samples, but *fastBE* found an increased amount of heterogeneity in the parent tumor and “CPT-11 Resistant” samples (Supplementary Figure 16).

## 6 Discussion

We defined a linear optimization problem, the 𝓁_1_-VAFRP, a subproblem of the NP-complete 𝓁_1_-VAFFP. By exploiting the special structure of the matrices which appear in this regression problem, we derived an algorithm which runs in 𝒪(*mnd*) time where *m* is the number of samples, *n* is the number of clones, and *d* is the depth of the input tree, 𝒯obtaining asymptotic and empirical speedups over state-of-the-art linear programming solvers. Using our regression algorithm, we developed a method *fastBE* for the 𝓁_1_-VAFFP which scales to large, multi-sample bulk DNA sequencing datasets. While *fastBE* serves as a practically useful tool for phylogenetic inference, we also believe our 𝓁_1_-regression algorithm and structured regression model is of independent interest, and will serve as a useful tool for the development of other algorithms for phylogenetic inference from multi-sample bulk DNA sequencing data.

There are several limitations of the present approach, which are directions for future work. On the theoretical side, it is an open question whether the time complexity of our regression algorithm can be improved from the current 𝒪 (*mnd*) time to the optimal 𝒪 (*mn*) time, which is the size of the input. On the practical side, extending our model and regression algorithm to additional classes of mutations evolutionary models is desirable. Here, we analyzed the simplest case of single nucleotide variants in copy neutral regions. Accounting for copy number heterogeneity by replacing the VAF with either the cancer cell fraction [47–49] or the descendant cell fraction [50], could improve the performance of our method on real datasets. Furthermore, using evolutionary models that allow for mutation loss – e.g., the Dollo model [5, 51], or generalizations [52, 53] – is a challenging future direction. Finally, extending *fastBE* to automatically infer mutation clusters, output uncertainty estimates, or enforce longitudinal consistency in temporal samples [10], may serve useful in expanding the scope and utility of *fastBE* to the broader scientific community.

## A Supplementary results

### A.1 An equivalent characterization of clonal matrices

We continue the study of clonal matrices by drawing an analogy to perfect phylogeny. Specifically, we introduce a new, recursive definition of clonal matrices that is inspired by the recursive definition of perfect phylogeny matrices described by Gusfield [28, 29]. Formally, we show that a matrix *B* is an *n*-clonal matrix if and only if the rows and columns of *B* can be reordered such that *B* is *clonally canonical*, a term we define below.

#### Definition 4.

*A matrix B is clonally canonical if i) B*_*i*,1_ = 1 *for all i* ∈[*n*], *ii) B*_1,*j*_ = 0 *for all j* ∈{2, …, *n*}, *and iii) there exists clonally canonical matrices B*_1_, *B*_2_, …, *B*_*k*_ *such that B*_2:*n*,2:*n*_ *is block diagonal with blocks B*_1_, *B*_2_, …, *B*_*k*_. *That is, B is clonally canonical if it has the form:*

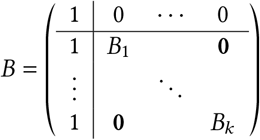

Conveniently, for a given clonal tree 𝒯, it is straightforward to construct a clonally canonical matrix associated with 𝒯 by relabeling the clones by their preorder traversal index during a depth first search starting at *r*. With this definition, we are now ready to state several equivalent characterizations of clonal matrices.

#### Proposition 3.

*For any binary n-by-n matrix B, the following conditions are equivalent:*

i. *B is an n-clonal matrix*.
ii. *The rows and columns of B can be reordered to make B clonally canonical*.
iii. *B satisfies the following conditions [5]:*
  a. *There exists exactly one index r such that* 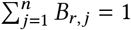.
  b. *For all i* ≠ *r, there exists exactly one j such that B* _*j,k*_ = 1 *implies B*_*i,k*_ = 1 *and* 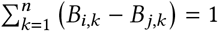.
  c. *B*_*i,i*_ = 1 *for all indices i* ∈ [*n*].

The clonally canonical form of the clonal matrix associated with a clonal tree 𝒯 is especially useful since it enables efficient multiplication and inversion of *B*.

#### Proposition 4.

*Let B be an n-by-n clonally canonical matrix associated with the clonal tree 𝒯. Then, the four matrix vector products*

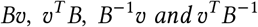

*can be computed in 𝒪* (*n*) *time for any vector* υ ∈ ℝ^*n*^ *when given the clonal tree 𝒯*.

## B Supplementary proofs

### Proposition 3.

*For any binary n-by-n matrix B, the following conditions are equivalent:*

i. *B is an n-clonal matrix*.
ii. *The rows and columns of B can be reordered to make B clonally canonical*.
iii. *B satisfies the following conditions [5]:*
  a. *There exists exactly one index r such that* 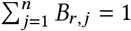.
  b. *For all i* ≠ *r, there exists exactly one j such that B* _*j,k*_ = 1 *implies B*_*i,k*_ = 1 *and*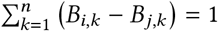.
  c. *B*_*i,i*_ = 1 *for all indices i* ∈ [*n*].

*Proof*. Suppose the theorem holds for all *k < n*. Clearly, the theorem holds for the base case when *n* = 1.

(*i*) ⇒ (*ii*) : Let 𝒯 be the *n*-clonal tree associated with *B*. Consider a relabeling of the clones of 𝒯 by their preorder traversal index during a depth first search of starting at the root. Then, *r*(𝒯) is assigned index 1 and the children of *r*(𝒯) are assigned indices *i*_1_ *<* … *< i*_*k*_ where *k* is the number of children of *r*(𝒯). Let *B*^′^ be the clonal matrix associated with 𝒯 under this relabeling. Then, 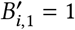 and 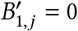 for all *j* ∈ {2, …, *n*} since all clones contains the mutation 1 and the root only contains the mutation 1. Let *B*_1_, …, *B*_*k*_ be defined recursively as the clonally canonical matrices associated with the subtrees rooted at the children *i*_1_, …, *i*_*k*_ of *r*(𝒯), which exist by the inductive hypothesis. Then, *B*^′^ is clonally canonical with blocks *B*_1_, …, *B*_*k*_.

To see this, observe that all vertices 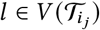 satisfy *i* _*j* −1_ *< l < i* _*j*+ 1_ as we performed a depth first traversal to relabel 𝒯. This implies that the sub-matrix *B*^′^ [*l*_*i*_ : *l*_*i* +1_ −1, *l*_*i*_ : *l*_*i* +1_ −1] is equal to *B*_*i*_, where we set *l*_*k* +1_ = *n* +1. Since *B*_*i*_ are clonally canonical, this completes this direction of the proof.

The equivalence of (*i*) and (*iii*) follows from the proof of Lemma 1 in [5]. It follows that (*ii*) implies (*i*) as clonally canonical matrices trivially satisfy (*iii*).

### Proposition 4.

*Let B be an n-by-n clonally canonical matrix associated with the clonal tree 𝒯*. *Then, the four matrix vector products*

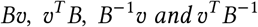

*can be computed in 𝒪*(*n*) *time for any vector* υ ∈ ℝ^*n*^ *when given the clonal tree 𝒯*.

*Proof*. By Lemma 1, the clonal matrix inverse *B*^−1^ = (*I* − *A*) and has exactly 2*n* non-zero entries. As such, we can compute the products *B*^−1^υ and υ^*T*^ *B*^−1^ in 𝒪 (*n*) time as the tree 𝒯 defines the indices of non-zero entries.

Next, consider a clonally canonical matrix *B* with clonally canonical blocks *B*_1_, …, *B*_*k*_ with dimensions *n*_1_, …, *n*_*k*_ such that 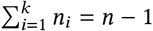. Define the coordinates 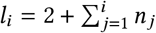 and *row vectors* 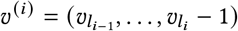, such that υ^*T*^ = (υ_1_, υ ^(1)^, …, υ ^(*k*)^). Then, by the properties of block matrix multiplication,

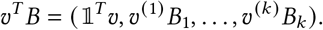

However, since *B*_1_, …, *B*_*k*_ are also clonally canonical, we have that the first entry of υ ^(*i*)^*B*_*i*_ is the sum of the entries of υ ^(*i*)^, implying that

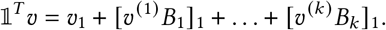

Therefore, after computing the *k* products υ ^(*i*)^*B*_*i*_, we can compute the υ^*T*^ *B* with only *k* + 1 additional operations. Assuming inductively that the number of operations to compute the product *u*^*T*^ *B*^′^ for any clonally canonical matrix *B*^′^ of size less than *n*^′^ *< n* is bounded by 2*n*^′^ − 1, the total number of operations to compute υ^*T*^ *B* is at most

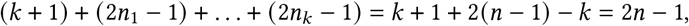

proving that υ^*T*^ *B* is computable in 𝒪 (*n*) time.

To see that the product *B*υ can also be efficiently computed, treat υ ^(*i*)^ as a *column* vector. Then, by the properties of block matrix multiplication,

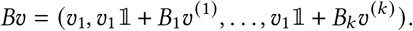

Therefore, after computing the *k* products *B*_*i*_υ ^(*i*)^ with an offset term υ_1_, we obtain *B*υ with only a single operation, proving that *B*υ is computable in 𝒪 (*n*) time by a similar argument as above.

### Lemma 1.

*For any clonal matrix B associated with a clonal tree* 𝒯 *having adjacency matrix A, B* = (*I* − *A*)^−1^. *Consequently*, [*B*^−1^υ]_*i*_ = υ_*i*_ − υ_*δ* (*i*)_ *if i* ≠ *r* (𝒯), *and* [*B*^−1^υ]_*i*_ = υ_*i*_ *when i* = *r* (𝒯).

*Proof*. Observe that 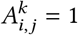 if and only if *j* is an ancestor of *i* in 𝒯 at tree distance *k* from *i*. Then,

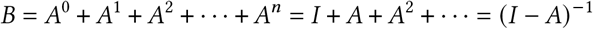

where the first equality follows from the observation above. The second equality from the observation that *A*^*k*^ is the zero matrix when *k > n*.

### Corollary 2.

*Let f be a frequency vector of length n, let B be an n-by-n clonal matrix, and let* 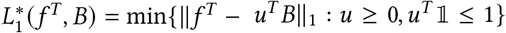. *Then* 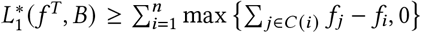, *or the total violation of the sum condition* (11). *Thus*, 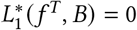 *if and only if the sum condition is satisfied*.

*Proof*. The first part of the corollary follows Lemma 1 of El-Kebir et al. [5]. The second part of the corollary follows from observing that

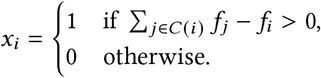

is a feasible solution to the DPTLP in Theorem 2.

## C Supplementary methods

### C.1 Simulation details

Our simulation procedure loosely follows the steps outlined in AncesTree [5]. Briefly, we simulate an *n*-clonal tree 𝒯, a *m*-by-*n* usage matrix *U*, and assign each of the *k* mutations to one of the *n* clones. Then, for each sample-mutation pair, we sample total and variant read counts.

In more detail, we constructed two sets of simulated instances, referred to as the *small* and *large* simulated instances respectively. The small simulated instance is defined by the following set of five parameters:

- *n* ∈ {3, 5, 10, 20, 30, 50}: the number of clones,
- *m* ∈ {5, 10, 25, 50}: the number of samples,
- *k* ∈ {500}: the number of mutations,
- *d* ∈ {40}: the read depth,
- *s* ∈ {1, …, 11, 12}: the random seed,

yielding a total of 288 small simulated instances. The large simulated instance is defined by the following set of parameters:

- *n* ∈ {100, 250, 500, 1000}: the number of clones,
- *m* ∈ {50, 100}: the number of samples,
- *k* ∈ {2000}: the number of mutations,
- *d* ∈ {40}: the read depth,
- *s* ∈ {0, 1, 2, 3, 4, 5}: the random seed,

yielding a total of 48 large simulated instances.

For a fixed set of parameters (*n, m, k, d, s*), we perform the following sequence of steps to create our simulation:

1. Randomly sample an *n*-clonal tree 𝒯with root vertex 0 from the set of all *n*-clonal trees using Wilson’s algorithm.
2. Randomly assign each of the *k* mutations to one of the *n* clones to obtain a (surjective) map from mutations to clones *ψ* : [*k*] → [*n*].
3. Randomly sample a usage matrix *U* row-by-row:
  a. For each row *i* ∈ [*m*], sample the number of non-zero entries *l*_*i*_ ∼ Uniform(1, …, *n*).
  b. For row *i* ∈[*m*], select the indices of the *l*_*i*_ non-zero usage proportions uniformly at random. Then, select the value of the *l*_*i*_ non-zero proportions from a Dirichlet distribution.
4. Set the ground-truth frequency matrix *F* = *U B*_𝒯_.
5. Sample the total read counts ^*T*^_*i,j*_ ∼ Poisson(*d*) for all samples *i* ∈ [*m*] and mutations *j* ∈ [*k*].
6. Sample the variant read counts *V*_*i,j*_ ∼ Binomial(^*T*^_*i,j*_, *F*_*i,ψ* (*j*)_) for all samples *i* ∈ [*m*] and mutations *j* ∈ [*k*].

### C.2 Running phylogenetic methods

To run *fastBE*, CITUP [7], CALDER [10], Pairtree [11], and Orchard [12] on the simulations, we passed in the variant read counts *V*, the total read counts ^*T*^, and the mutation to clone mapping *ψ* : [*k*] →[*n*]. For *fastBE* and CALDER [10], we first estimated the frequency matrix 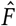 as

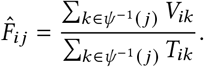

When running CITUP [7], we used the quadratic integer programming version of the method and set the max jobs equal to the number of compute cores. For CALDER, we used the 𝓁_1_ objective and default parameters. For Pairtree [11], we used the default parameters and explicitly set the number of threads equal to the number of available compute cores. Since Pairtree outputs a posterior distribution over trees rather than a single tree, we select the most likely tree output by their tool. For Orchard [12], we used the recommended settings outlined in their Github repository. Similarily, we select the most likely tree output by their tool.

To benchmark the regression algorithm described in (Section 3.3) against the linear programming based approaches implemented by the solvers Gurobi [43] and CPLEX [44], we constructed larger simulated instances with *n* ∈ {250, 500, 750, 1000} clones, *m* ∈ { 50, 100, 200, 500} samples, and *k* ∈ {500, 1000, 2000, 4000} mutations at a read depth of *d* = 50. For each of the three regression algorithms, we passed in the ground truth frequency matrix *F* and clonal matrix *B*_𝒯_, and then measured both the wall-clock and (internal) solver time to infer the usage matrix *U* which minimizes the 𝓁_1_ error ∥*F*−*UB*_𝒯_ ∥_1_. All simulations and method evaluations were performed on the computing cluster provided by the Princeton University Department of Computer Science. Each simulated instance and method evaluation was performed independently on cluster nodes with 8 GB of memory and 16 compute cores.

### C.3 Evaluation metrics

Our evaluation procedure is philisophically similar to [5–7, 9, 10] in that we believe there is a single ground truth clone tree and usage matrix, and that the goal of factorization is to recover this ground truth. In contrast, the evaluation philosophy of Pairtree [11] believes that the inherent ambiguity leads to many, equally plausible clone trees and usage matrices, and that the goal of factorization is to learn this set (or distribution) of clone trees and usage matrices. Therefore, we evaluate methods with respect to their deviation from the ground truth clonal tree 𝒯, usage matrix *U*, and frequency matrix *F*.

We first measure the deviation between the inferred clonal tree 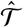 and the ground truth clonal tree 𝒯. To perform this measurement, we compute the set of ancestral relationships for each tree, and measure how similar these sets are to each other. Formally, we define the set of ancestral relationships as follows,

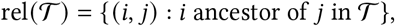

where we say (*i, j*) is a *positive* if (*i, j*) ∈ rel (𝒯)and a *negative* otherwise. Then, the total number of positive and negatives is,

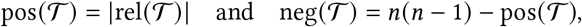

and the false positive and false negative rate of 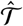 predicting positive and negatives ancestral relationships is,

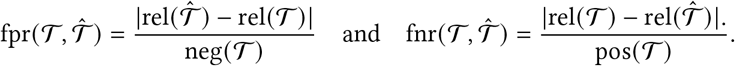

Similarly, we define precision, recall, and the F1-score, which is the harmonic mean of precision and recall.

We next measure the deviation between the inferred and ground truth usage and frequency matrices. In particular, given an estimated usage matrix Û and frequency matrix 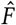, the matrix error is the normalized 𝓁_1_ error:

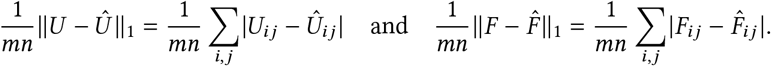

When methods do not explicitly report Û and 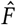, and instead only output 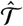, to compute the matrix error we first need to estimate Û and 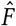.To perform this estimation, we conservatively set Û to be the minimizer of 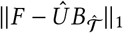 over all usage matrices Û and then set 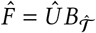. This estimation procedure ensures the matrix error 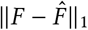 is minimized for the reported clonal tree.

### C.4 fastBE details

While we outlined our stochastic search algorithm *fastBE* in Section 4, we omitted a formal description of our procedure for choosing an order to add mutations and the efficient recomputation approach. First, to choose an order in which to add mutations, we use the *F-sum trick* introduced in Orchard [12]. In particular, we use the order *O* = { *o*_1_, …, *o*_*n*_} obtained by sorting the column sums of *F* in descending order. That is, the *F*-sum trick orders the mutations *O* such that,

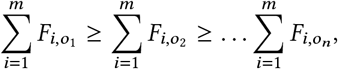

the intuition being that mutations with higher frequency should occur earlier in the evolutionary process. Second, by the latter part of Theorem 1, after the addition of a mutation to 𝒯, it takes 𝒪 (*mnd*) time to recompute the score *L*^*^ (*F, B* _𝒯_). Further, the adoption of a child of the parent can be implemented as an SPR move, also taking only 𝒪 (*mnd*) time to recompute the score *L*^*^ (*F*,*B*_𝒯_) upon. Performing adoptions in the correct order then leads to the total time complexity of 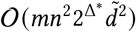 over all possible placements.

When applying *fastBE* to clusters of mutations, as specified by a map *ψ* : [*k*] → [*n*] from mutation to cluster identity, we define the input frequency matrix 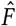 as

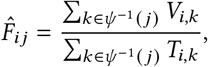

where *V*_*i,j*_ and ^*T*^_*i,j*_ is the variant and total read count of mutation *j* in sample *i*. That is, the columns of 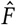 correspond to the average frequency of the mutations in each cluster.

## List of Supplementary Figures

**Supplementary Figure 1:**
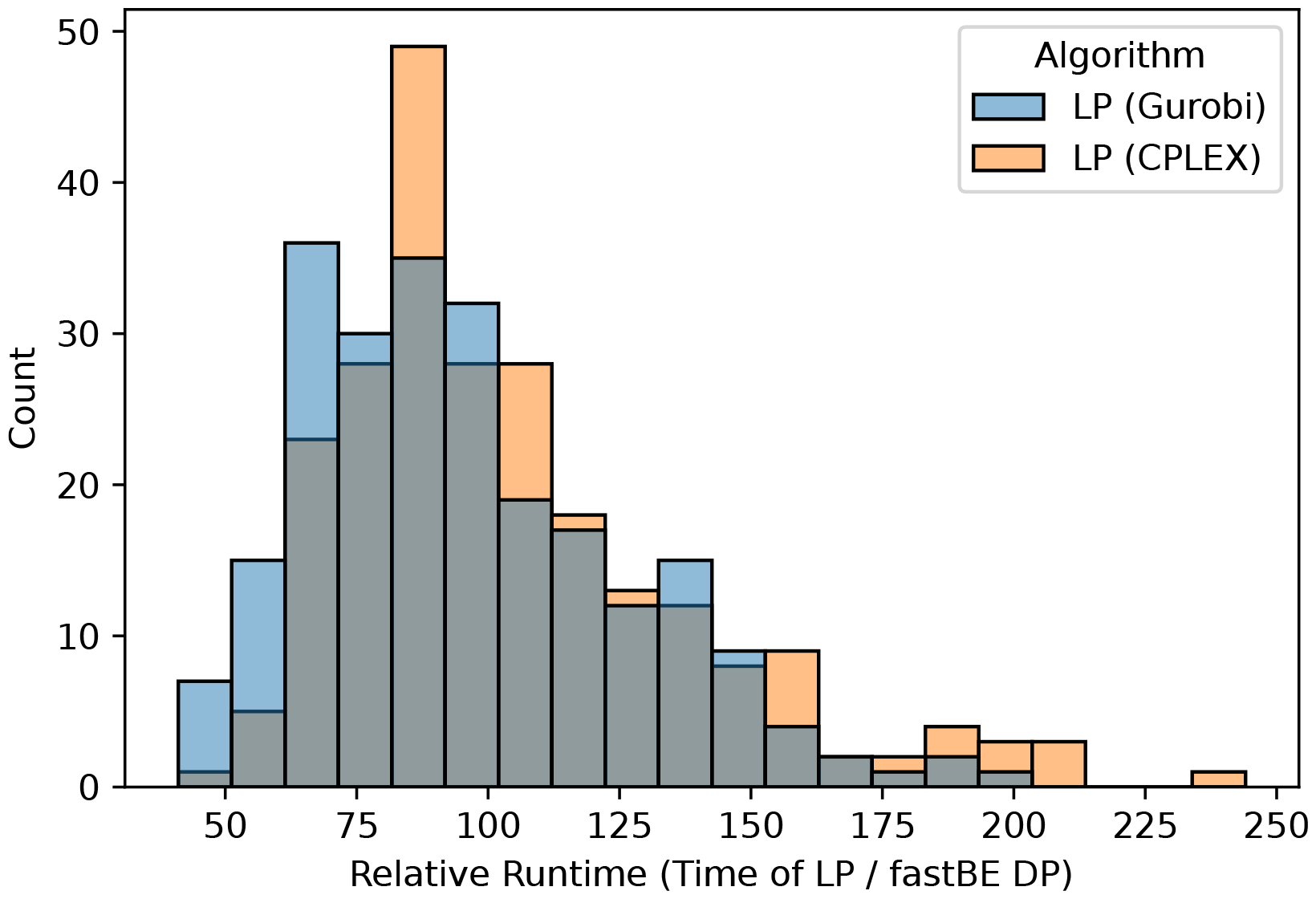
Relative wall-clock runtime comparison of our algorithm to that of two commercial linear programming solvers, Gurobi and CPLEX. Only the time required to solve the model is reported.

**Supplementary Figure 2:**
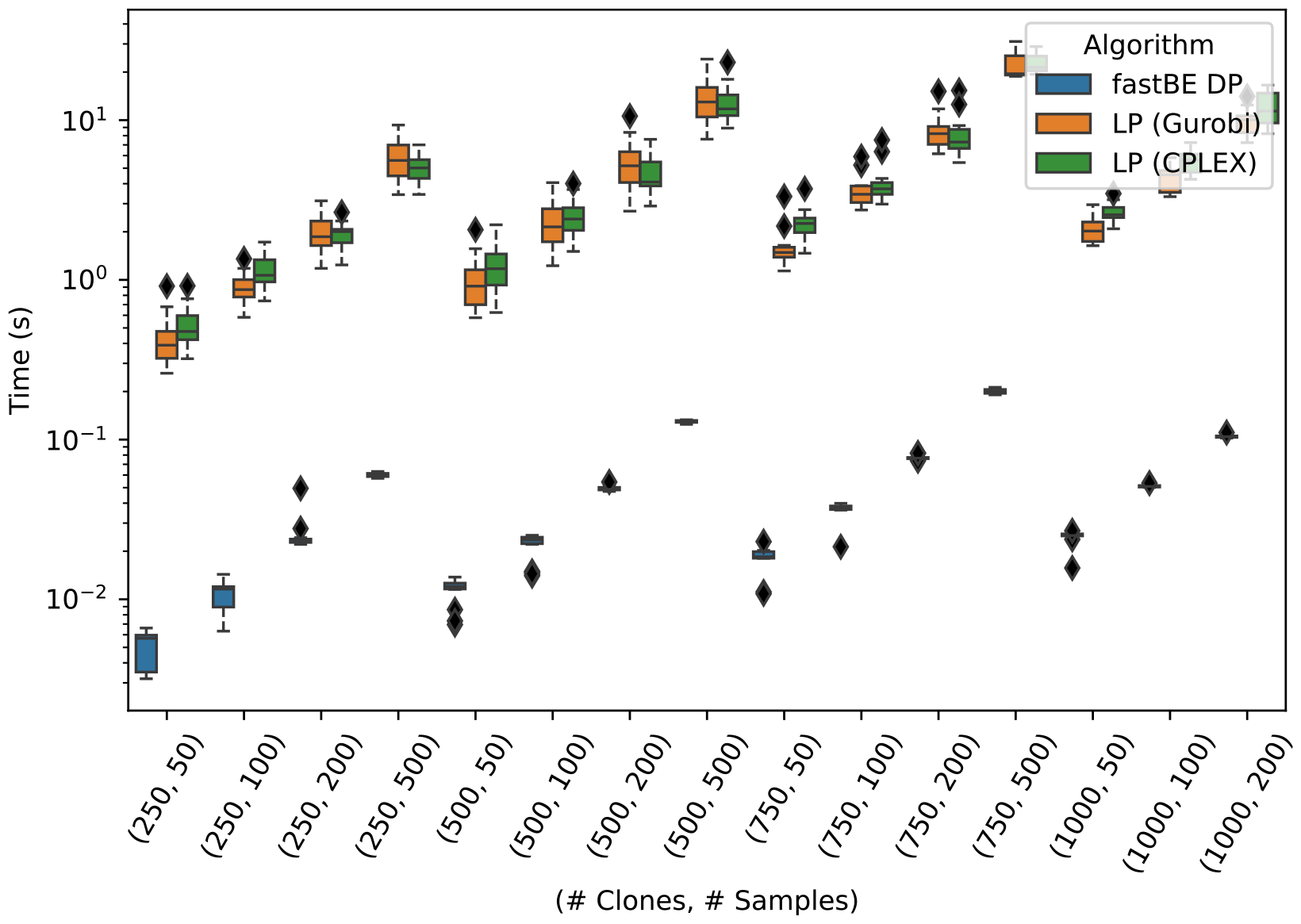
Absolute wall-clock runtime comparison of our algorithm to that of two commercial linear programming solvers, Gurobi and CPLEX for a varying numbers of clones *n* = 250, 500, 750, 1000 and samples *m* = 50, 100, 200, 500. Only the time required to solve the model is reported.

**Supplementary Figure 3:**
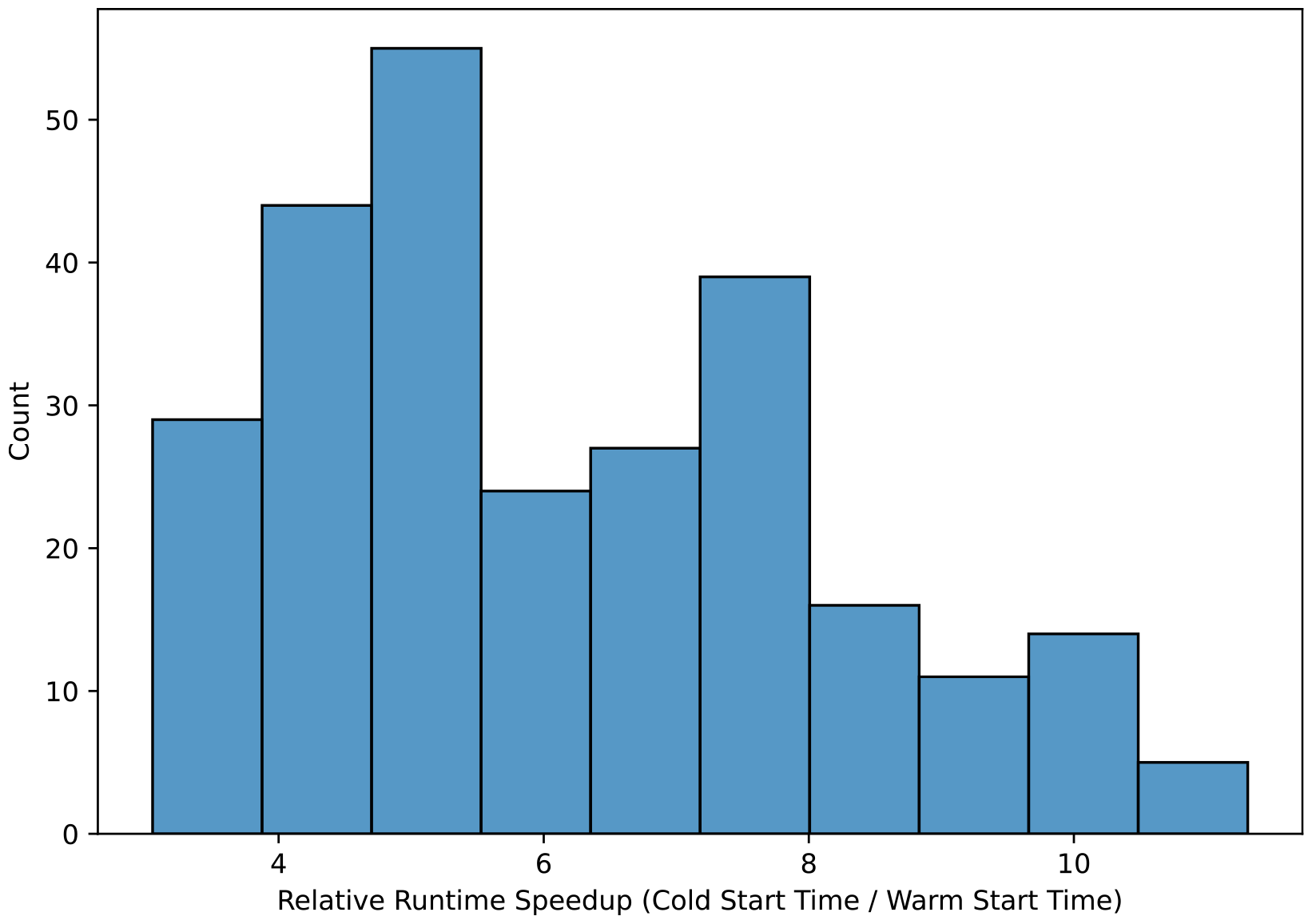
Relative wall-clock runtime comparison of warm versus cold starting our regression algorithm across 264 instances with varying numbers of clones *n* = 250, 500, 750, 100 and samples *m* = 50, 100, 200, 500.

**Supplementary Figure 4:**
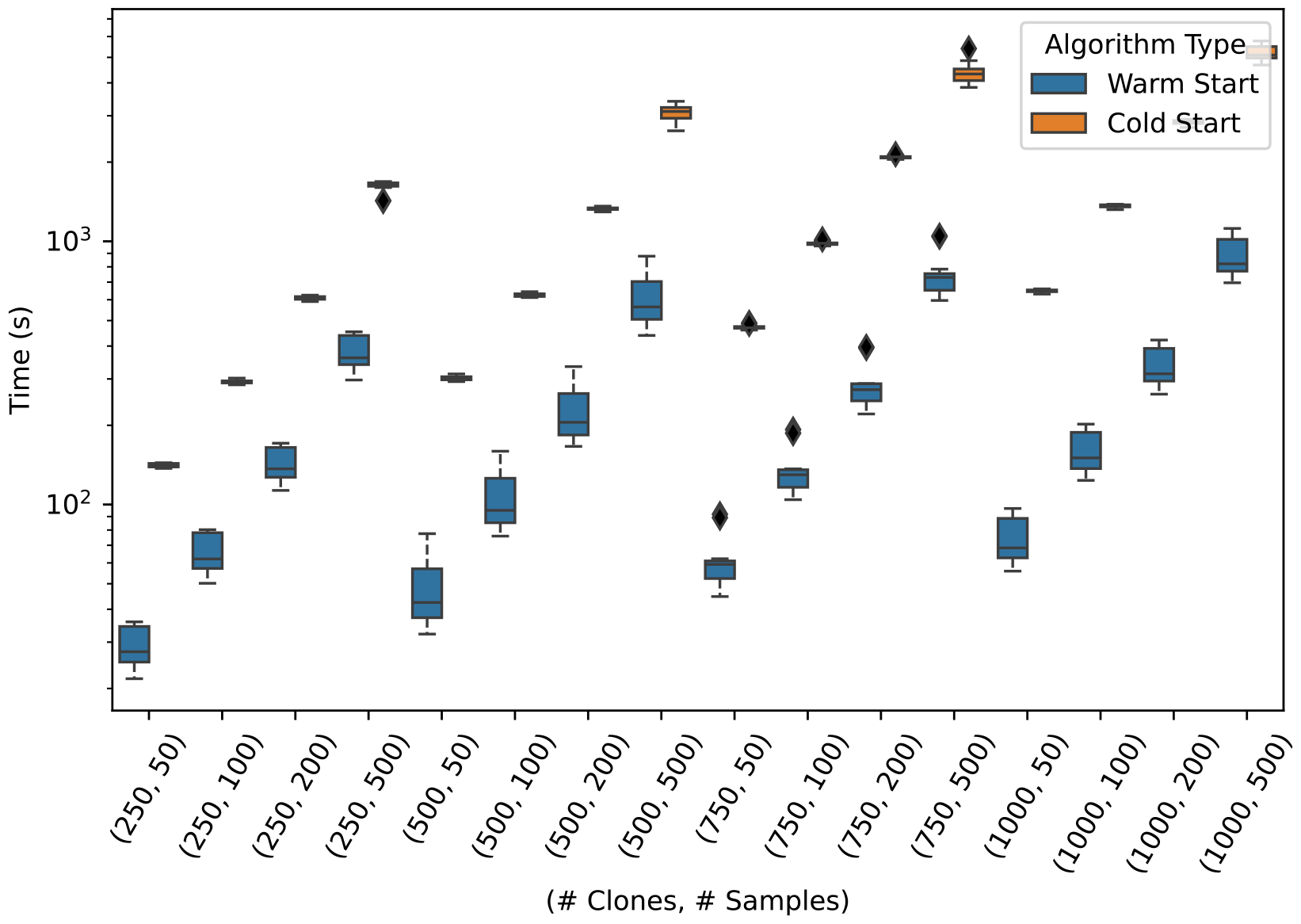
Absolute wall-clock runtime comparison of warm and cold starting our regression algorithm across 264 instances with varying numbers of clones *n* = 250, 500, 750, 100 and samples *m* = 50, 100, 200, 500.

**Supplementary Figure 5:**
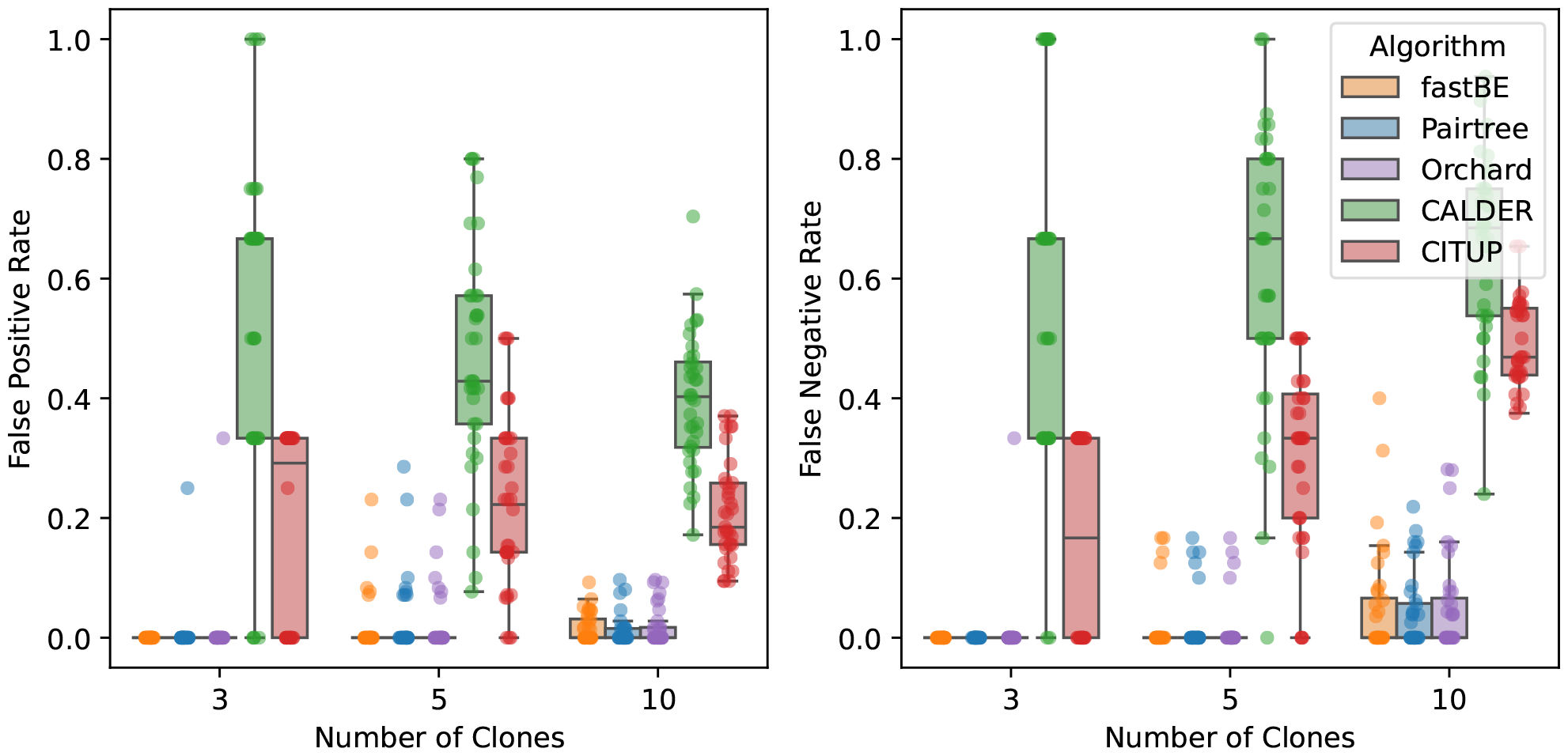
The **(left)** false positive rate and **(right)** false negative rate of reconstructing pairwise relationships for *fastBE*, Pairtree [11], Orchard [12], CALDER [10], and CITUP [7] on simulated data for small instances with ≤ 10 clones and ≤ 25 samples.

**Supplementary Figure 6:**
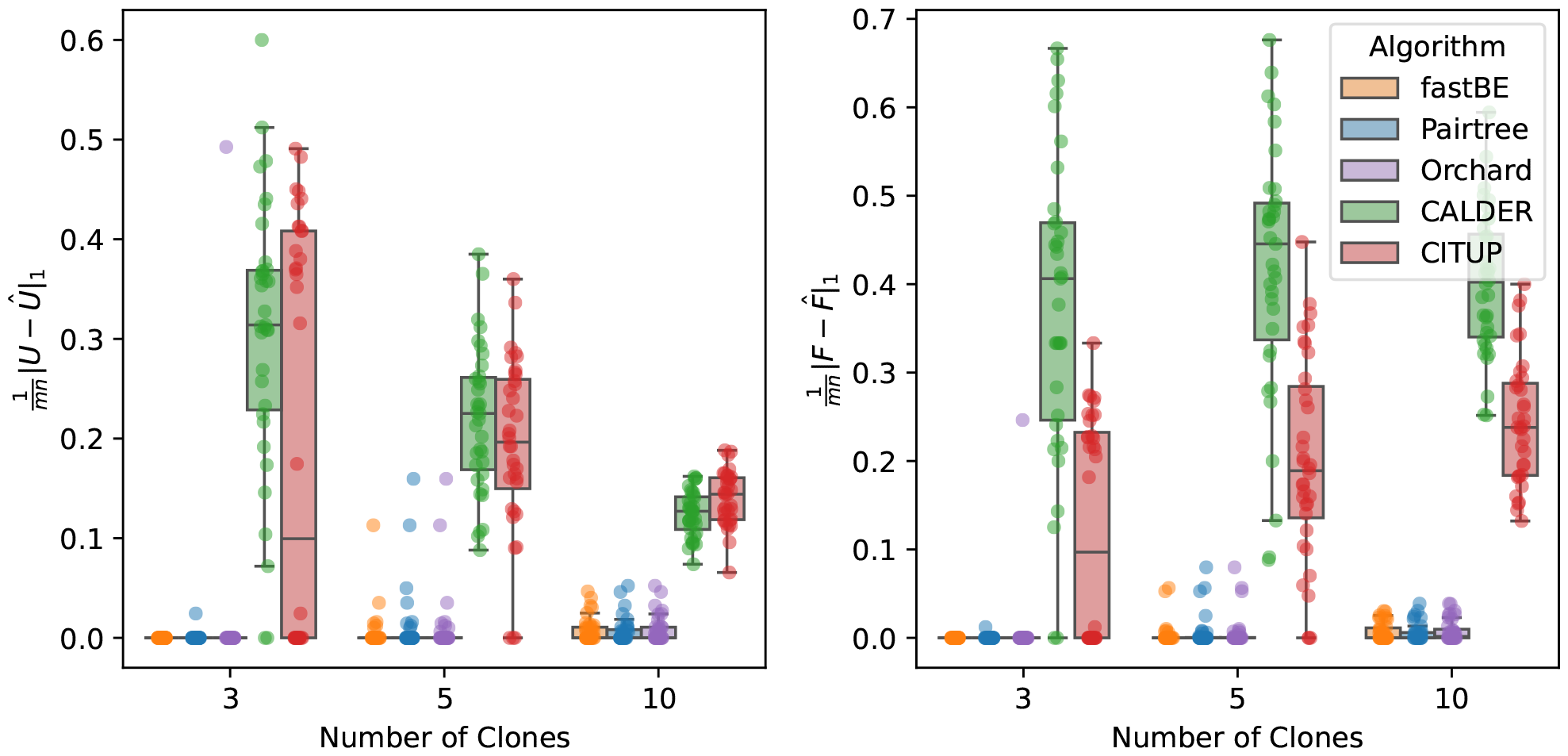
The normalized 𝓁_1_ matrix error **(left)** ∥*U* − Û ∥_1_ and **(right)** 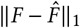 for *fastBE*, Pairtree [11], Orchard [12], CALDER [10], and CITUP [7] on simulated data for small instances with ≤ 10 clones and ≤ 25 samples.

**Supplementary Figure 7:**
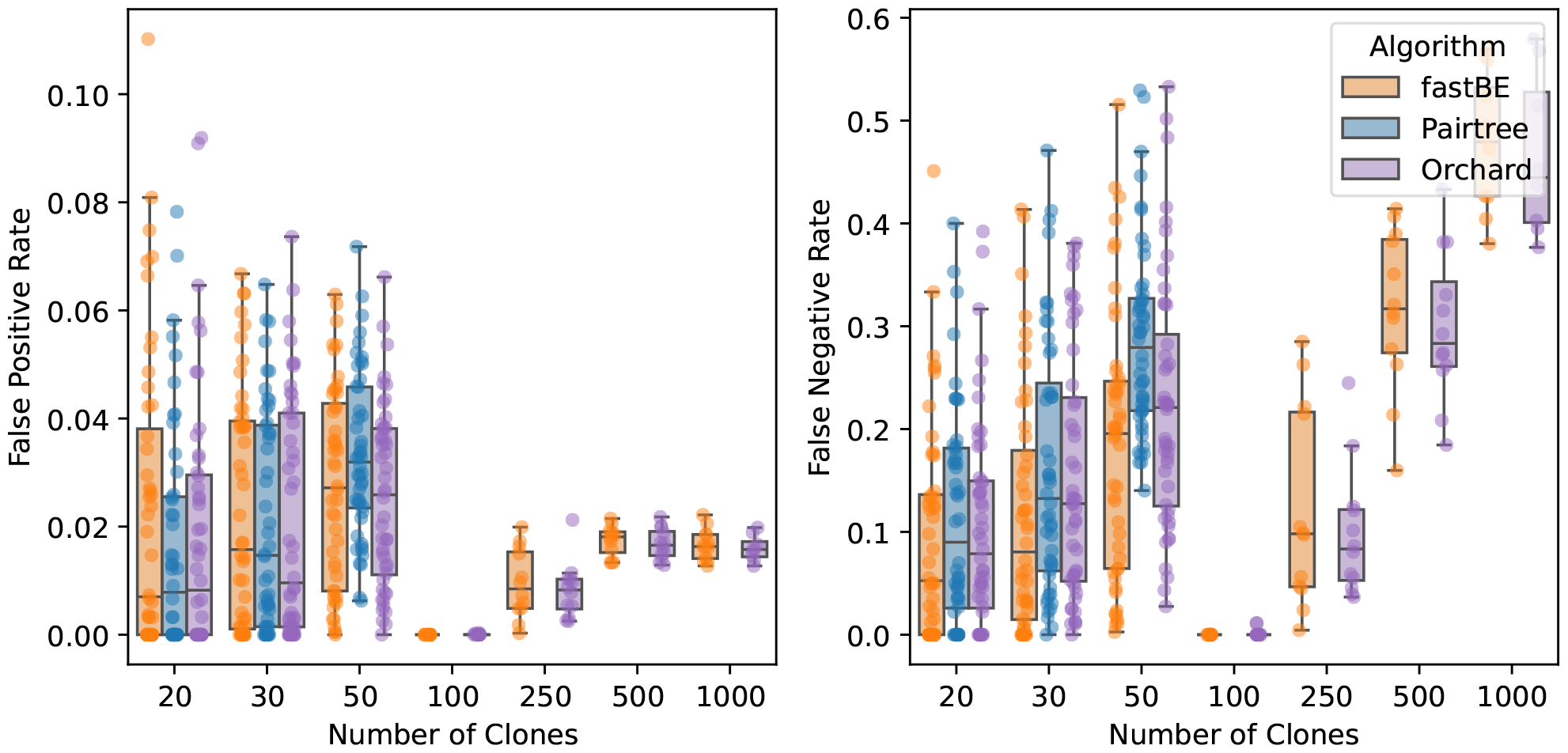
The **(left)** false positive rate and **(right)** false negative rate of reconstructing pairwise relationships for *fastBE*, Pairtree [11], and Orchard [12] on simulated data with ≥20 clones. Pairtree did not scale to the ≥100 clone setting and was excluded.

**Supplementary Figure 8:**
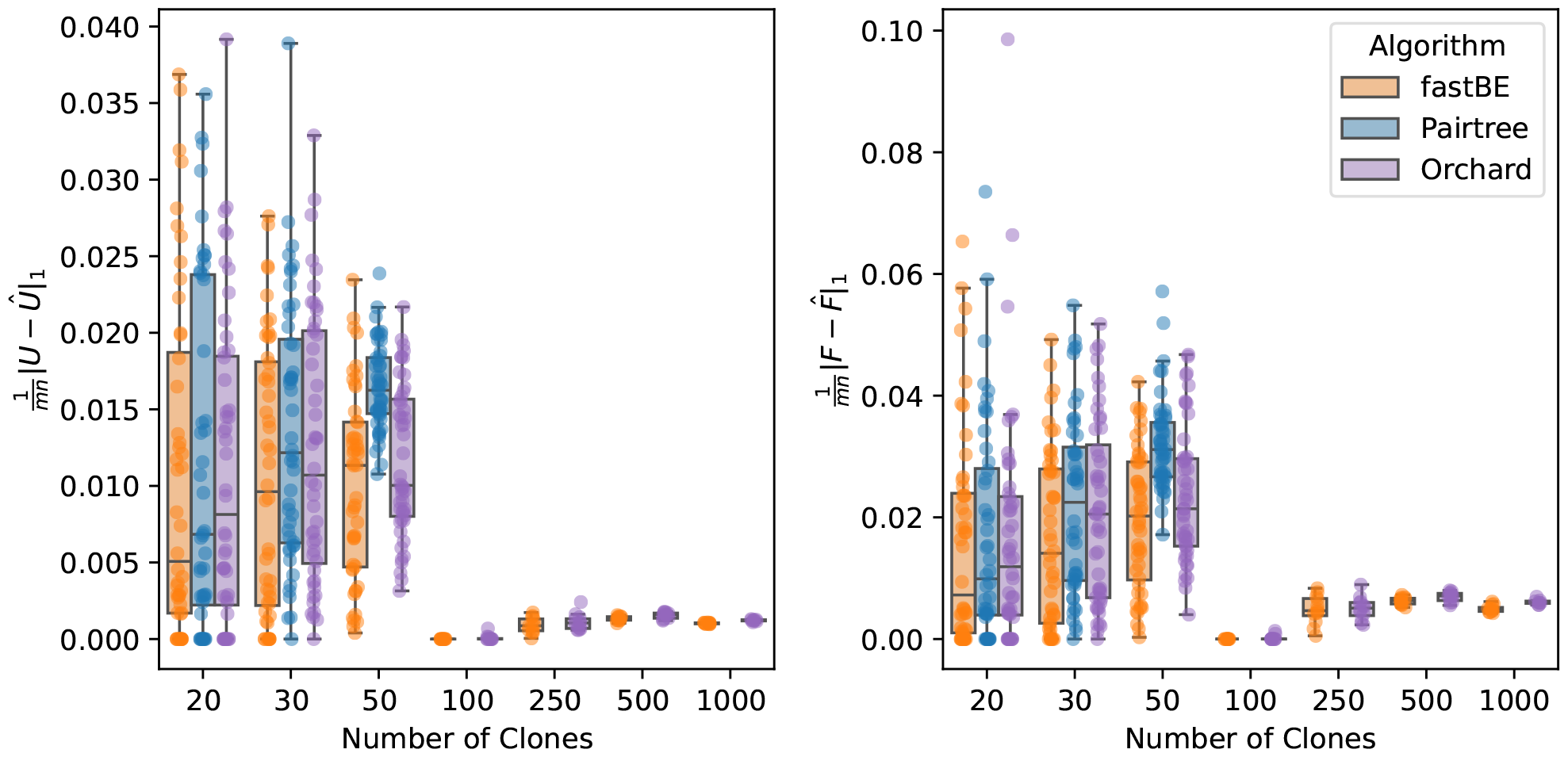
The normalized 𝓁_1_ matrix error **(left)** ∥*U* − Û ∥_1_ and **(right)** 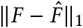 for *fastBE*, Pairtree [11], and Orchard [12] on simulated data with ≥ 20 clones. Pairtree did not scale to the ≥ 100 clone setting and was excluded.

**Supplementary Figure 9:**
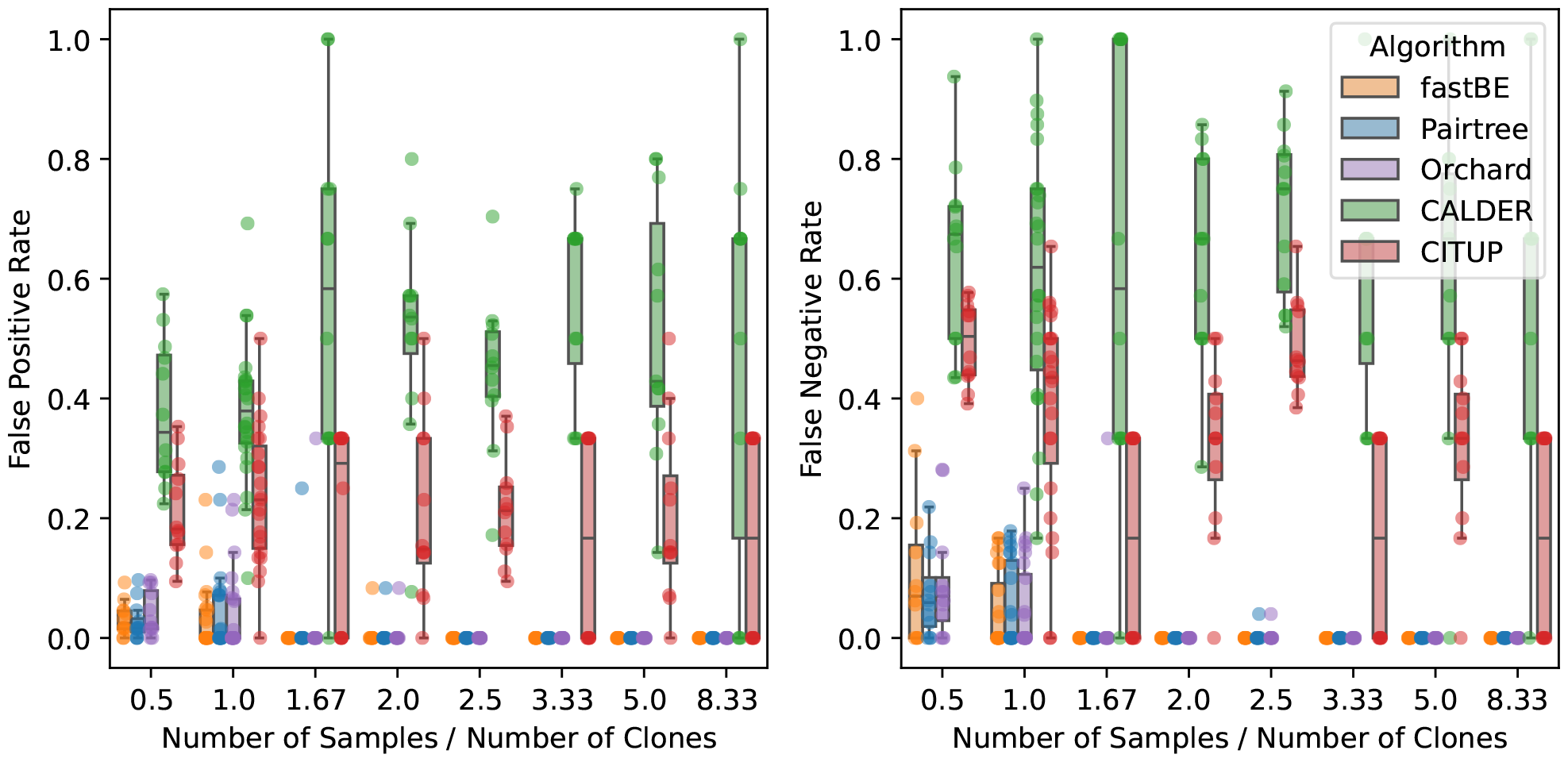
The **(left)** false positive rate and **(right)** false negative rate of reconstructing pairwise relationships for *fastBE*, Pairtree [11], Orchard [12], CALDER [10], and CITUP [7] on simulated data with ≤10 clones and ≤25 samples versus the ratio of samples to clones.

**Supplementary Figure 10:**
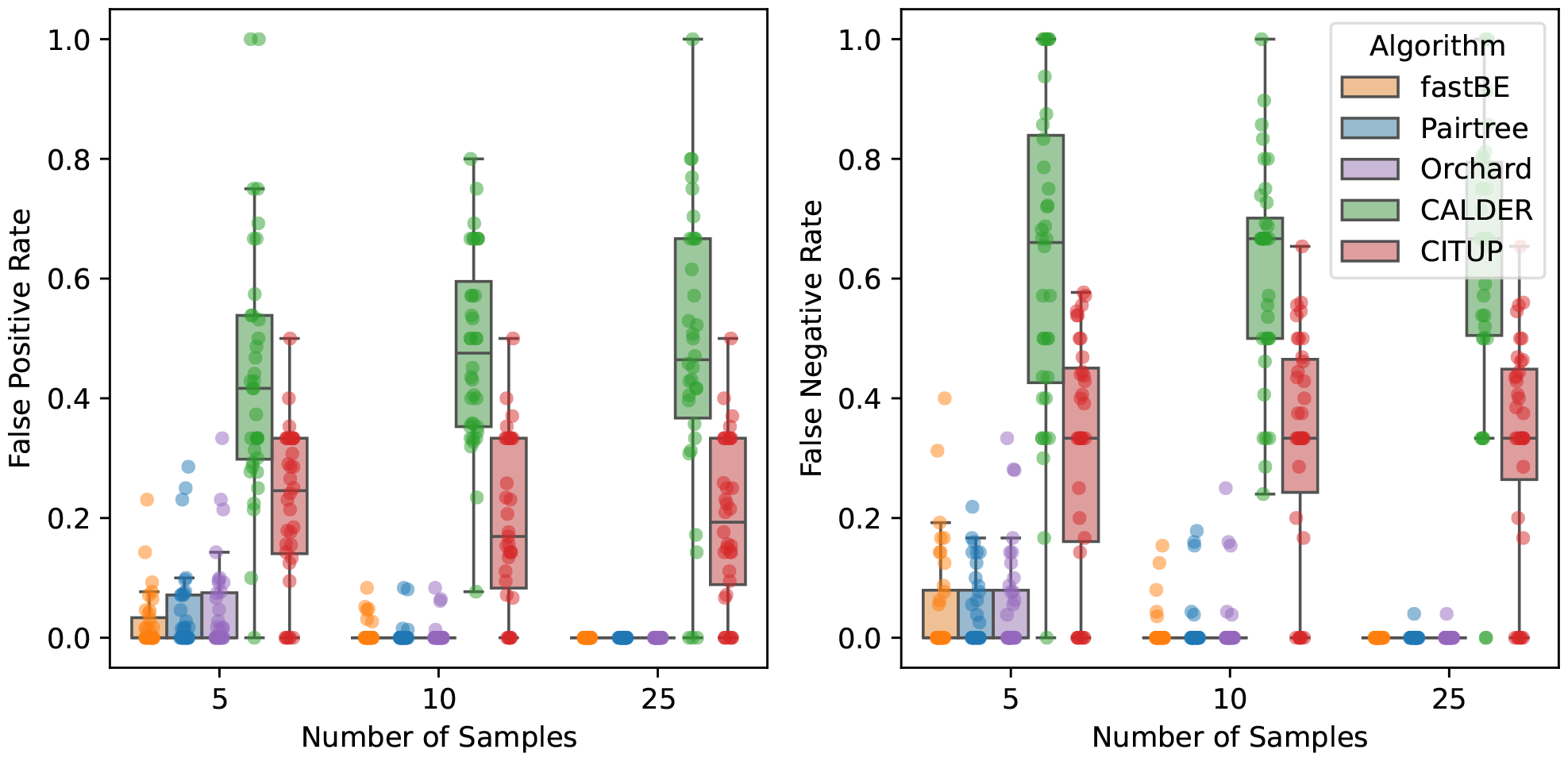
The **(left)** false positive rate and **(right)** false negative rate of reconstructing pairwise relationships for *fastBE*, Pairtree [11], Orchard [12], CALDER [10], and CITUP [7] for simulated data with ≤10 clones versus the number of samples.

**Supplementary Figure 11:**
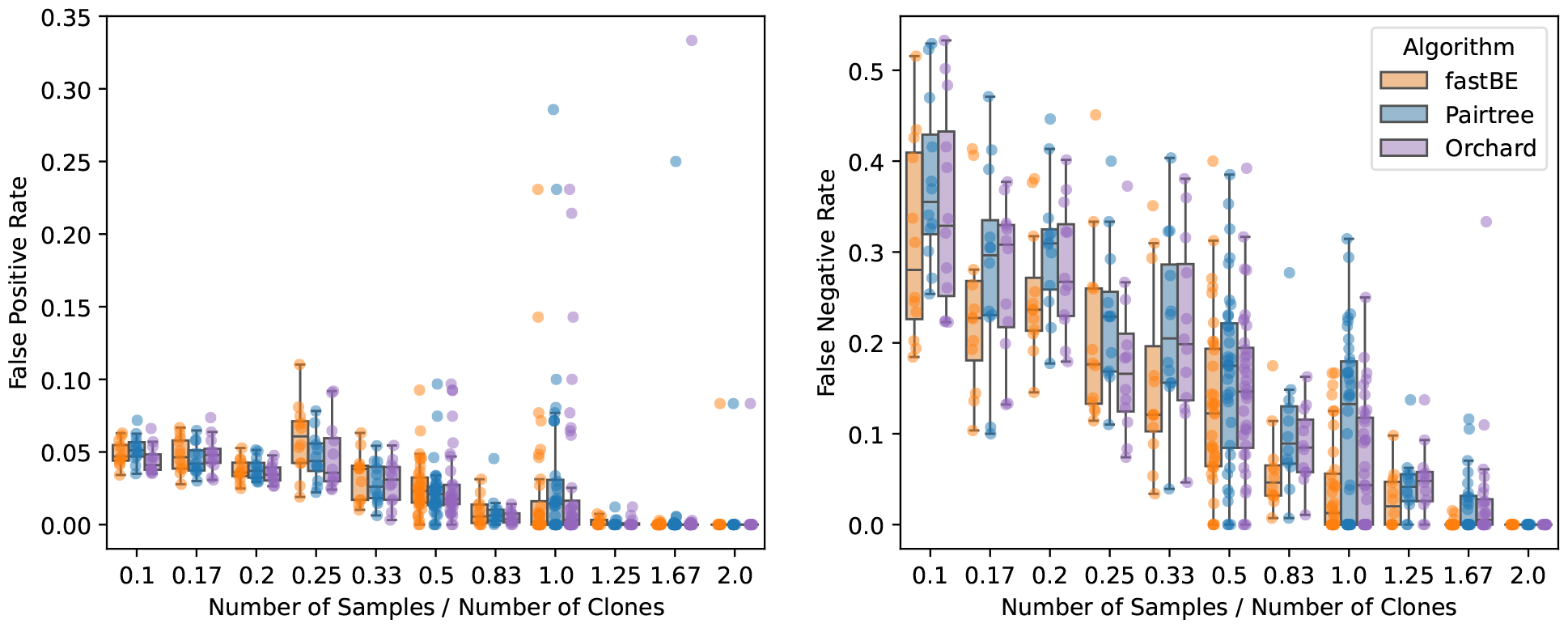
The **(left)** false positive rate and **(right)** false negative rate of reconstructing pairwise relationships for *fastBE*, Pairtree [11], and Orchard [12] on simulated data with ≥20 and ≤100 clones versus the ratio of samples to clones. When the ratio of samples to clones exceeded 2, all methods correctly recovered the ground truth tree and were excluded from the plots.

**Supplementary Figure 12:**
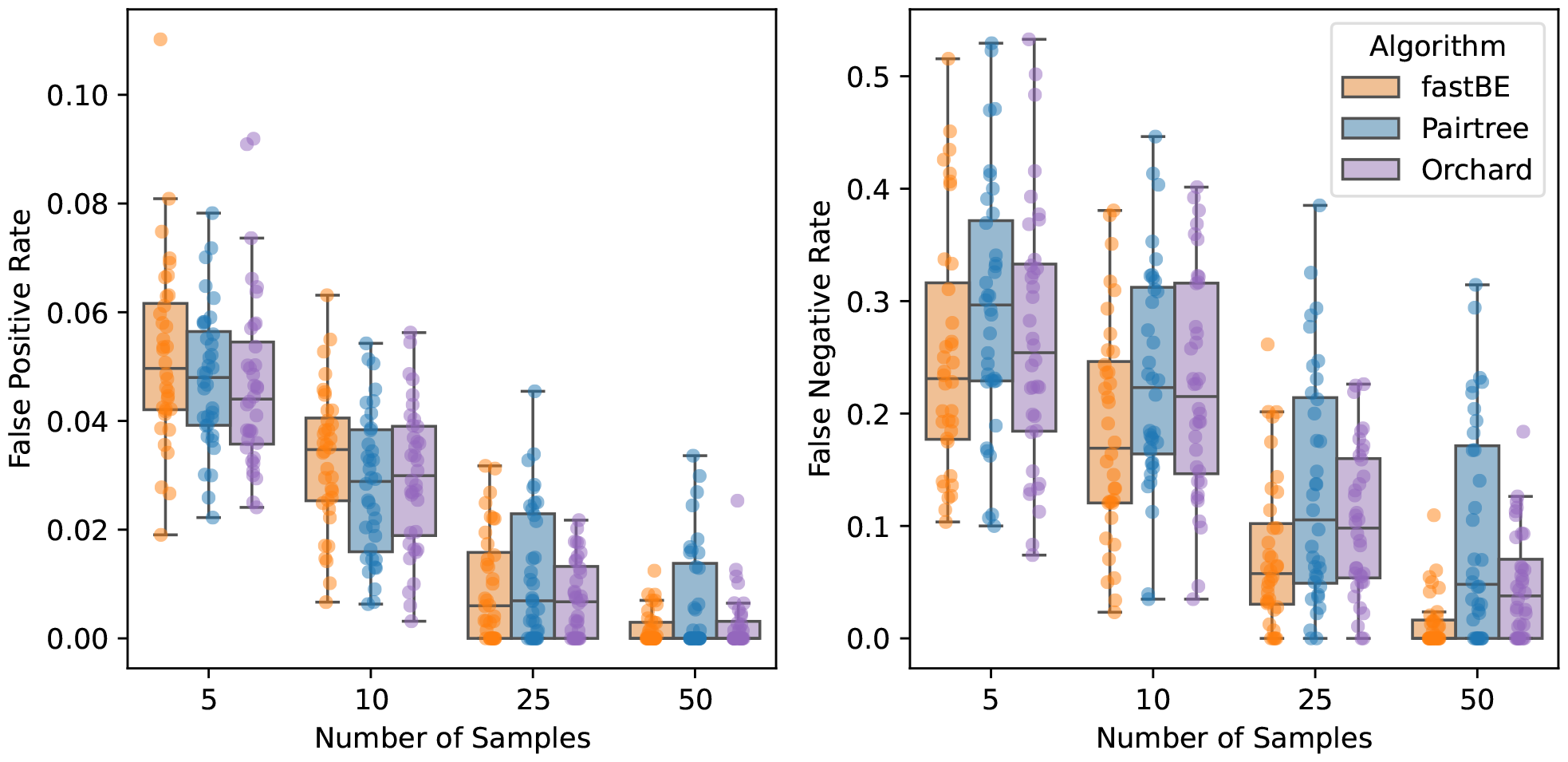
The **(left)** false positive rate and **(right)** false negative rate of reconstructing pairwise relationships for *fastBE*, Pairtree [11], and Orchard [12] on simulated data with ≥ 20 and ≤ 100 clones versus the number of samples.

**Supplementary Figure 13:**
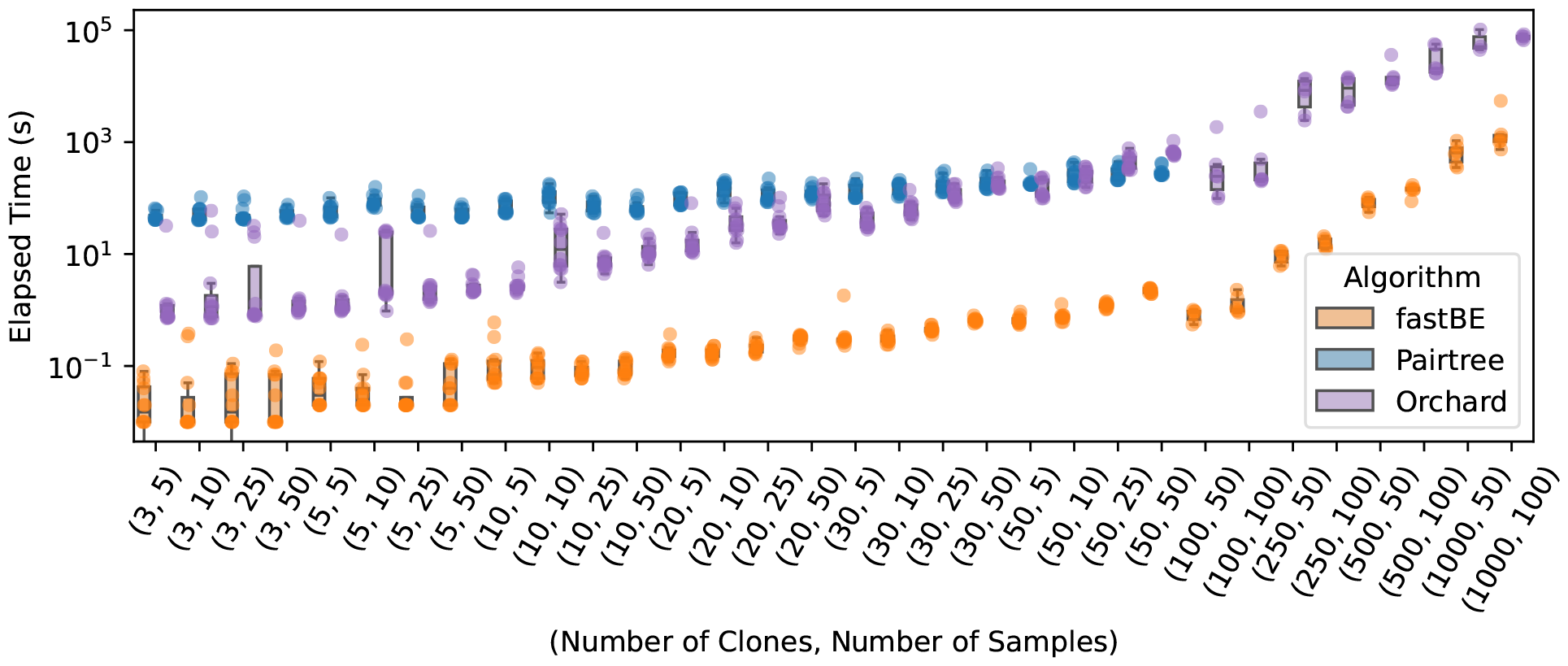
Absolute wall-clock runtime comparison of *fastBE*, Pairtree [11], and Orchard [12] with varying numbers of clones and samples. Pairtree did not scale to the ≥ 100 clone setting and was excluded.

**Supplementary Figure 14:**
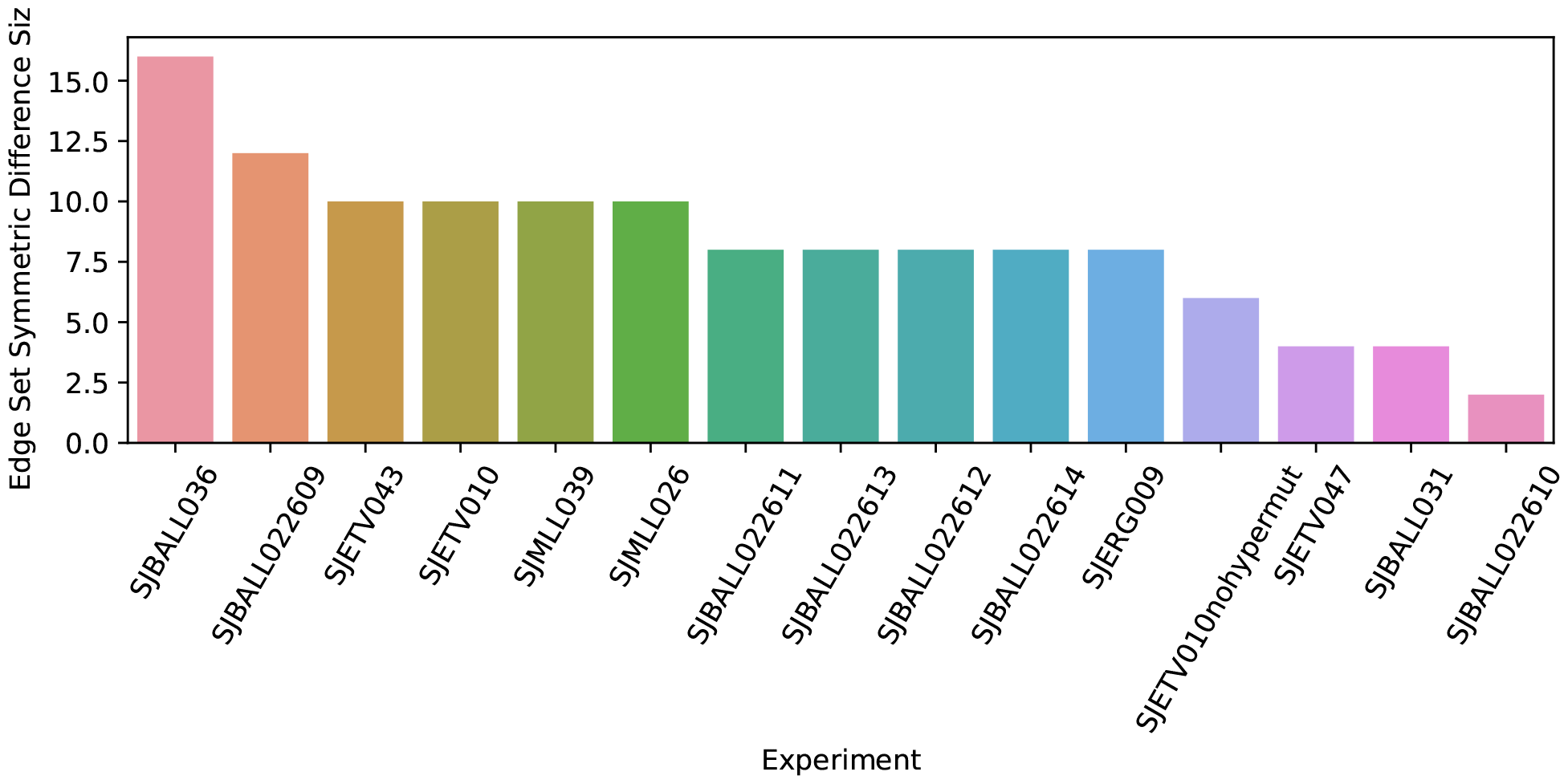
The size of the symmetric difference in the edge sets for the phylogenies inferred by *fastBE* and Pairtree [11] on data from fourteen patients with B progenitor acute lymphoblastic leukemia.

**Supplementary Figure 15:**
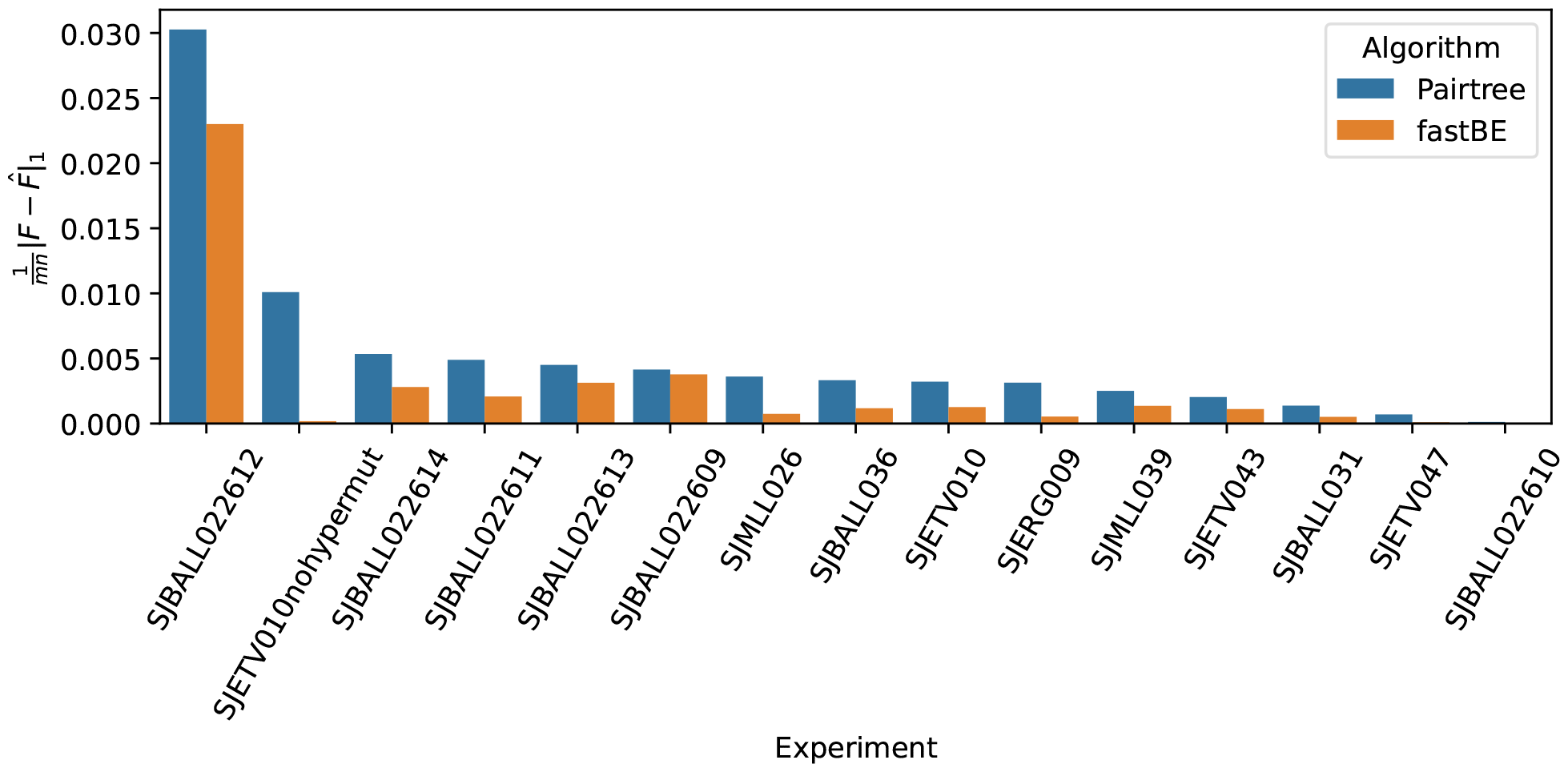
The normalized 𝓁_1_ matrix error for the phylogenies inferred by *fastBE* and Pairtree [11] on data from fourteen patients with B progenitor acute lymphoblastic leukemia.

**Supplementary Figure 16:**
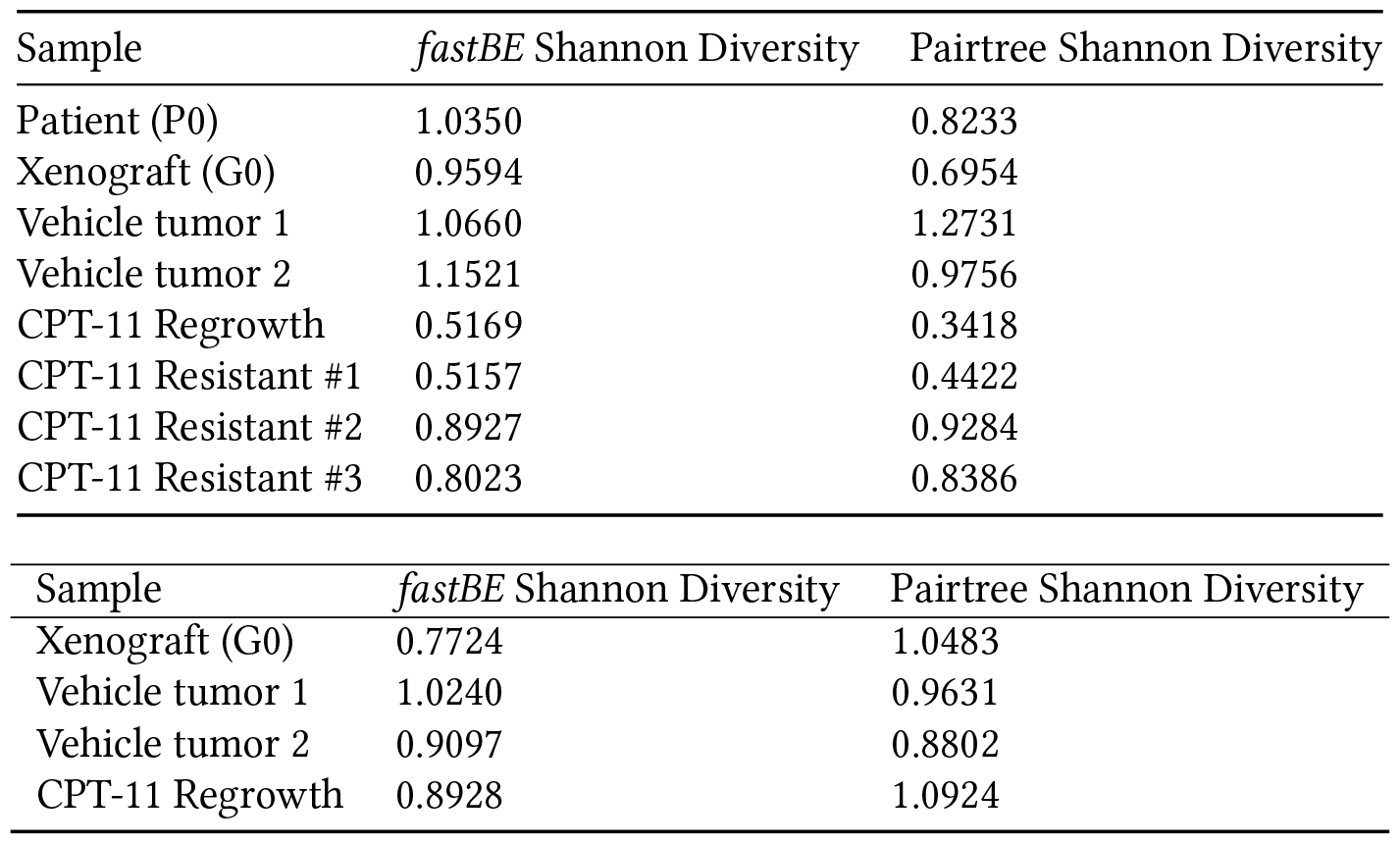
The Shannon diversity index estimates for phylogenies inferred by *fastBE* and Pairtree [11] on multi-sample bulk DNA sequencing performed on patient-derived colorectal cancer models **(top)** POP66 and **(bottom)** CSC28.

**Supplementary Figure 17:**
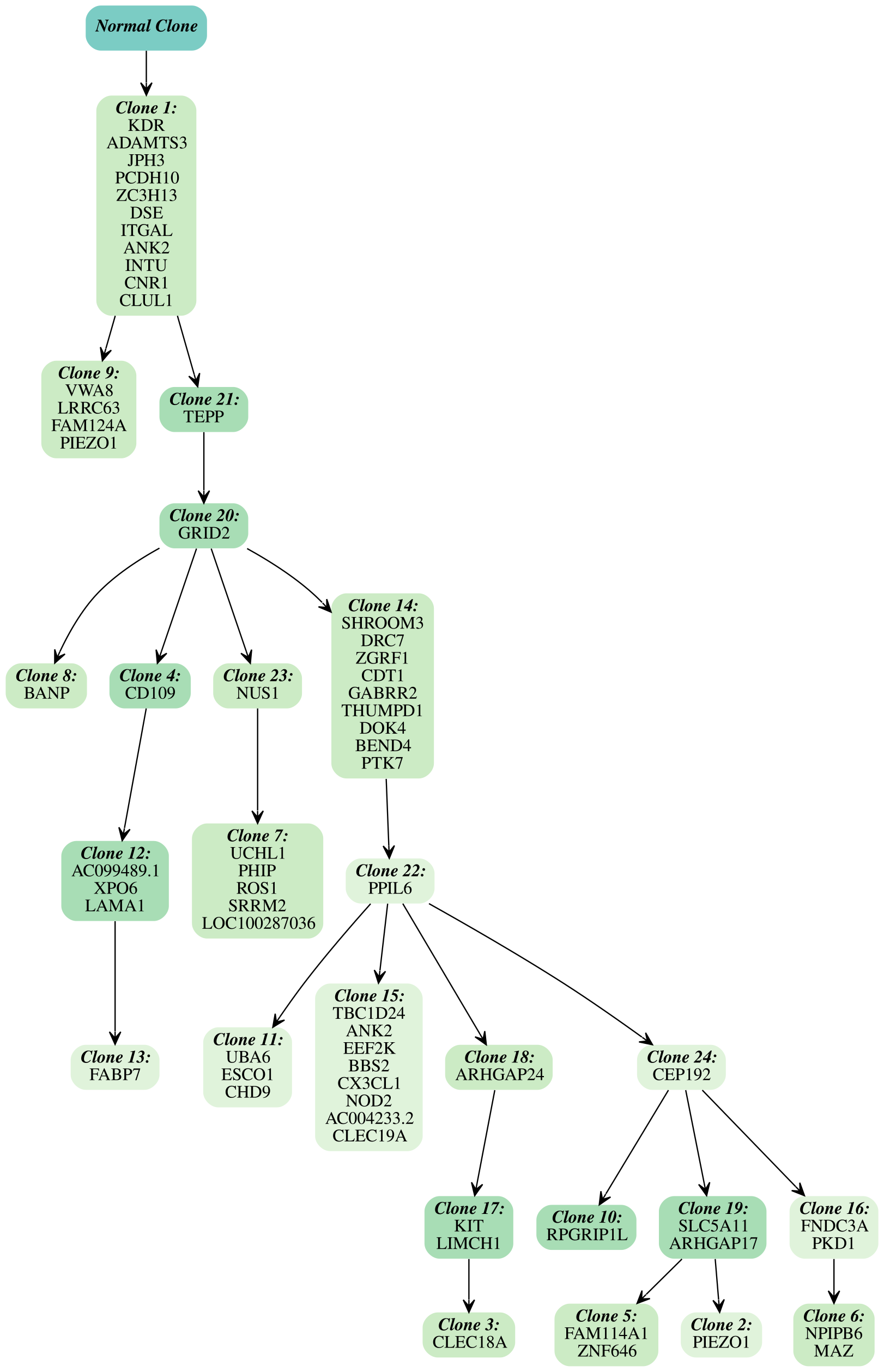
*fastBE* inferred phylogeny for the multi-sample bulk DNA sequencing performed on the CSC28 colorectal cancer model. Each vertex corresponds to a distinct clone and is labeled by the set of mutations which occur on the edge directed into the clone.

**Supplementary Figure 18:**
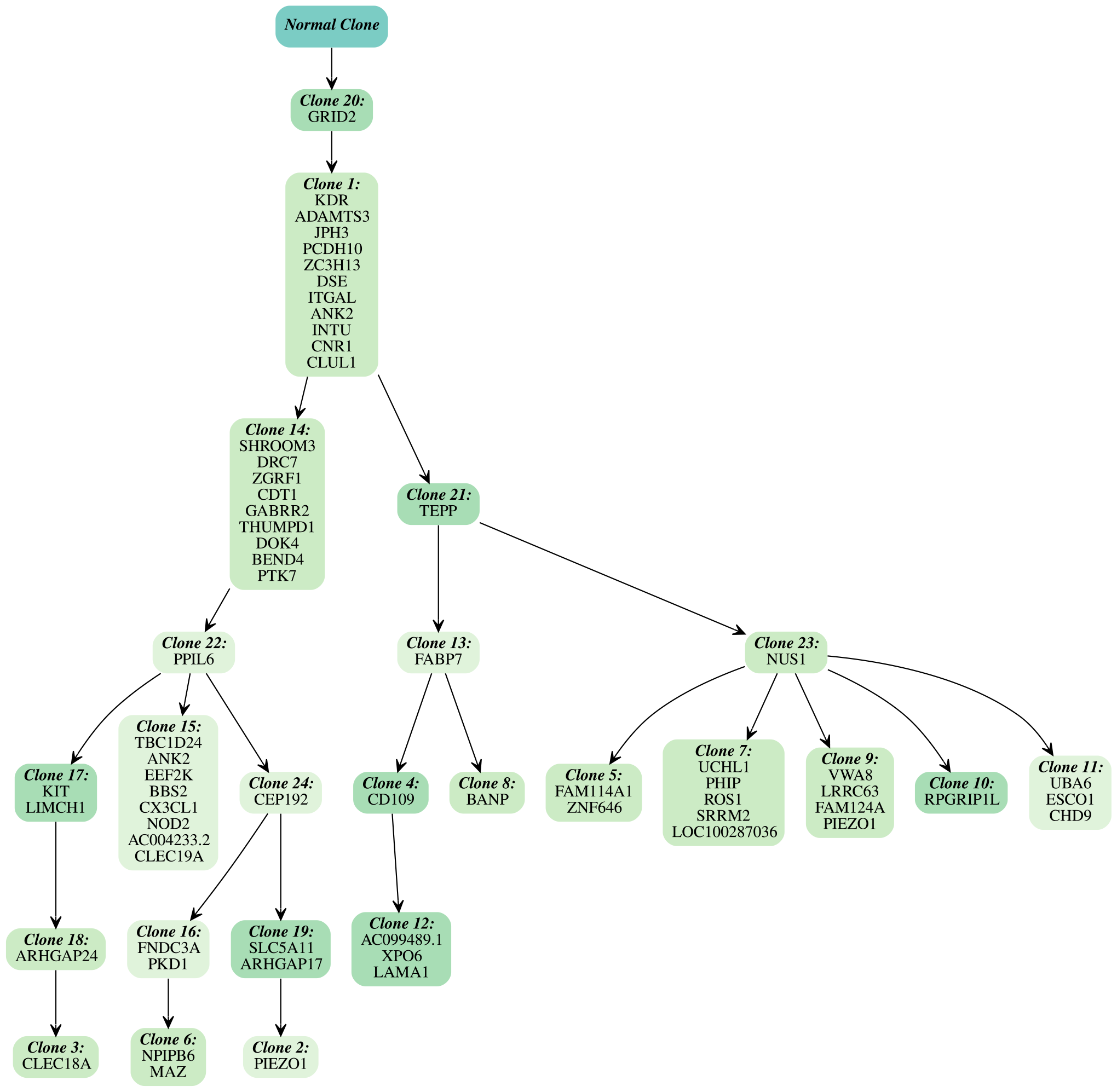
Pairtree inferred phylogeny for the multi-sample bulk DNA sequencing performed on the CSC28 colorectal cancer model. Each vertex corresponds to a distinct clone and is labeled by the set of mutations which occur on the edge directed into the clone.

## Footnotes

1 In practice, rather than modeling the evolution of individual mutations, mutations are typically clustered into mutation clusters using tools such as PyClone [25, 26] or SciClone [27], to make inference computationally tractable. These clusters of mutations are then assumed to both evolve together and satisfy the infinite sites assumption.

2 Our simplified definition of an *n*-clonal tree is equivalent to El-Kebir et al. [5] up to a renaming of the vertices in 𝒯.

